# Zone Matcher: A climate-based web application for deployment and assisted migration of forest trees

**DOI:** 10.1101/2025.03.30.644615

**Authors:** Tal J. Shalev, Meridith L. McClure, Nikolas Stevenson-Molnar, Gregory A. O’Neill, Tongli Wang, Joseph A. E. Stewart, Glenn T. Howe

## Abstract

Populations of forest trees are generally adapted to the climates they inhabit. The farther trees are moved from their local climates, the more long-term growth and survival tend to decrease. Current tree deployment and assisted migration rely on ‘climate distance thresholds’ (CDTs), which are climatic distances beyond which tree performance is considered unacceptable. Fixed zone systems, which have been used to guide deployment of native or orchard seedlots for more than 50 years, usually consist of contiguous geographic areas (zone units) divided into elevational bands (zones). In contrast, focal zone systems allow seed transfer among fixed zones that have similar climates. By using recent historical climates and future climate projections, focal zones can be used for current tree deployment or assisted migration. We developed a focal-zone system for the Pacific Northwest region of North America. First, we worked with stakeholders to select the base zones for the system. These consisted of geographic zone sets from Washington, Oregon, California, and Idaho/western Montana, and ecological zone sets from the U.S. and British Columbia. Second, by analyzing climate variation across the region, we developed a normalized Euclidean climate distance function consisting of nine climate variables from ClimateNA. Third, we inferred CDTs from analyses of climate variation within the base zones and from provenance tests. Fourth, we compared seed deployment areas using the fixed zone versus focal zone system, with and without assisted migration. Finally, we developed the Zone Matcher web application which implements our focal zone system. Across the region, we identified climate matches among 4,393 partially overlapping zones covering approximately 252 M ha. The unique area covered by these zones was about 167 M ha. Compared to fixed zones, our focal zone system increased the deployment area about 17-to 35-fold for the ecological zones and 70-to 300-fold for the geographic zones. This expands seed deployment options, allows more seedlots to be considered for a planting site, facilitates assisted migration, and simplifies sharing of seedlots among organizations. In addition to climate, seed transfer should also consider factors such as plantation soils, microtopography, and projections of competing vegetation, insects, diseases, and fire.

## INTRODUCTION

Nearly 100 years ago, foresters began to call for regulation of seed movement as the effects of inappropriate seed deployment became evident (Thrupp 1927, Bates 1929). Where non-local seed sources were used, forest trees often suffered from slower growth and lower survival (Isaac 1949, Langlet 1971). To improve the success of forest plantations, provenance tests were established as early as 1912 to understand the performance and adaptive variation of seed sources (Johnson et al. 2004, St.Clair et al. 2019). Local adaptation of trees—superior fitness of local versus non-local populations—is now broadly recognized in the ecological genetics literature of wide-ranging tree species (Carter 1996, Savolainen et al. 2007, Alberto et al. 2013). Despite some exceptions (Leimu and Fischer 2008, Candido-Ribeiro and Aitken 2024), local adaptation has given rise to the ‘local is best’ paradigm for seed source selection in reforestation and conservation plantings.

Decades of provenance tests confirmed that slower growth, lower survival, and greater pest damage often occur when seed sources are moved long climatic distances (Rehfeldt et al. 1999, Wilhelmi et al. 2017, Ahrens et al. 2019), and similar effects are expected from climate change (Rehfeldt et al. 2001, Wang et al. 2006, St.Clair and Howe 2007, Ikeda et al. 2014). Because trees often serve as foundation species, these effects can have negative cascading impacts on ecosystem function and diversity (Ellison et al. 2005, Ikeda et al. 2014). Planted forests, which occupy 293 million ha globally (FAO 2020), are key components of global biodiversity and other important ecosystem services (Millennium Ecosystem Assessment 2005). The use of climatically adapted trees will become increasingly important for ensuring planted forests remain healthy and productive (Hereford 2009).

In the United States, the ‘local is best’ paradigm gave rise to ‘seed transfer guidelines’ that were published as part of the USDA’s Forest Seed Policy of 1939. These guidelines suggested that seed should be planted (deployed) no more than 100 miles and 1,000 feet in elevation from its source location (Buck et al. 1970). In 1974, Campbell proposed using latitudinal and elevational ‘transfer distances’ to guide seed deployment (Campbell 1974). Seed transfer guidelines were subsequently refined based on new genecological information from major tree species (e.g., Campbell 1991, Sorensen 1992) and by using GIS technology. Seed transfer guidelines treat each seedlot or stand as being genetically distinct with its own deployment area. In contrast, because of substantial gene flow, most forest trees have large, more-or-less genetically homogeneous populations. Thus, the location-specific precision of these seed transfer guidelines is probably unnecessary. They also complicate the integration of seed inventories and reforestation planning, and are challenging to use with seedlots collected from many stands or from seed orchards. The use of seed transfer guidelines has also been called ‘floating’ and ‘focal point’ seed transfer (Rehfeldt 1983, Parker 1992, Ying and Yanchuk 2006).

To simplify seed deployment, ‘fixed’ seed zones began to be used in the 1960s, and are still widely used worldwide (Buck et al. 1970, Randall 1996, Randall and Berrang 2002, Ying and Yanchuk 2006, Ontario Ministry of Natural Resources and Forestry 2010). Fixed seed zones are geographical areas with fixed boundaries, within which local seed can be moved freely with low risk of maladaptation (Johnson et al. 2004, Ying and Yanchuk 2006). Like seed transfer guidelines, fixed zones keep seedlots collected from wild stands from being moved outside their zone of origin. Fixed seed zones guide the deployment of seed from native stands, whereas breeding zones guide the deployment of seed from breeding programs (e.g., seed from seed orchards, Howe et al. 2006). Thus, we use ‘deployment zone’ or ‘zone’ to refer to the deployment areas associated with either seed zones or breeding zones.

In 1966, the Western Forest Tree Seed Council (WFTSC) published a fixed zone system for Oregon and Washington. Zone units consisting of contiguous areas were subdivided into non-contiguous elevational bands that were used as seed zones. Because there were few genetic tests to rely on, the WFTSC used geography, climate, and vegetation to infer patterns of adaptive genetic variation (Western Forest Tree Seed Council 1972). In general, the intent has been to delineate fixed zones using the environmental factors responsible for genetic differentiation within species (Campbell 1986, Ying and Yanchuk 2006, Hamann et al. 2011, Potter and Hargrove 2012, Bower et al. 2014, Orquera et al. 2024).

In the Pacific Northwest region of North America, seed zones were first designed for Douglas-fir (*Pseudotsuga menziesii* (Mirb.) Franco), and later for other commercial tree species (Randall 1996, Randall and Berrang 2002, Johnson et al. 2004, Ying and Yanchuk 2006). Fixed zones make it easy to manage seed inventories and reforestation activities; thus, they were widely adopted. Over time, a variety of fixed zone sets (many of which overlap) were developed by different states, provinces, federal agencies, and private organizations (Howe et al. 2006). These include zone sets delineated primarily using geographic variables as climate surrogates (geographic zone sets) and others delineated primarily using ecological criteria (ecological zone sets) (Table 1, Appendix S1: Table S1).

**Table 1.** Tree deployment zones in the Pacific Northwest.

| **(a) General characteristics of geographic and ecological zone sets** |  |  |  |  |  |
| --- | --- | --- | --- | --- | --- |
| Type of zone set | Zone units |  | Zones |  |  |
|  | Type of zone units* | Delineation of zone units<sup>†</sup> | Type of Zones* | Delineation of zones<sup>†</sup> | Zone sizes |
| Geographic | Contiguous geographic areas | Climate (M)<br>Soils and geology (L)<br>Vegetation (M) | Non-contiguous elevation bands | Climate (H)<br>Soils and geology (M)<br>Vegetation (M) | Smaller |
| Ecological | No zone units *per se* (see zones) | No zone units *per se* (see zones) | Non-contiguous ecoregions | Climate (H)<br>Soils and geology (H)<br>Vegetation (H) | Larger |
| **(b) Specific characteristics of the Pacific Northwest zone sets** |  |  |  |  |  |
| Abbreviation | Type of zone unit<sup>‡</sup> | Coverage | Organization<sup>§</sup> | Year | Species<sup>¶</sup> |
| ***Geographic zone sets*** |  |  |  |  |  |
| CA | Topographic | CA | USFS | 1970 | Generic |
| ID/MT | Gridded | ID and MT (partial) | IETIC | 2020 | Generic |
| OR66 | Topographic | OR | WFTSC | 1966 | Generic |
| OR96 | Topographic | OR | ODF | 1996 | Species-specific |
| WA02 | Topographic | WA | WDNR | 2002 | Species-specific |
| WA66 | Topographic | WA | WFTSC | 1966 | Generic |
| ***Ecological zone sets*** |  |  |  |  |  |
| BEC | BEC subzone variants | BC | BCMof | 2021 | Generic |
| EPA4 | EPA Level IV ecoregions | US | EPA | 1986 | Generic |
| *Because the ecological deployment zones have no hierarchy, the delineated areas (zones) are analogous to both the zone units and zones of the geographic zone sets.<br><sup>†</sup>Letters indicate ability to account for variation in climate, soils and geography, or vegetation among adjacent zone units or zones (L = low, M = moderate, H = high).<br><sup>‡</sup>Topographic units are based on topographic features, gridded units are based on a 0.5-degree geographic grid, and BEC and EPA4 units are based on ecosystem classification.<br><sup>§</sup>BCMof is British Columbia Ministry of Forests, EPA is Environmental Protection Agency, IETIC is Inland Empire Tree Improvement Cooperative, ODF is Oregon Department of Forestry, USFS is United States Forest Service, WFTSC is Western Forest Tree Seed Council, and WDNR is Washington Department of Natural Resources.<br><sup>¶</sup>Generic zones are designed to be used for any species. OR96 and WA02 have 22 and 16 zone sets for individual species, respectively, but we analyzed only 10 species-specific zone sets that had existing shapefiles. |  |  |  |  |  |

The advent of GIS technologies led to renewed interest in seed transfer guidelines. Parker and van Niejenhuis, for example, combined geographic or climatic data and GIS technology to implement seed transfer guidelines in jack pine (*Pinus banksiana* Lamb.) and black spruce (*Picea mariana* (Mill.) B.S.P.), an approach they called ‘focal point’ seed deployment (Parker 1992, Parker and van Niejenhuis 1996). First, they used climate variables to predict patterns of genetic variation in adaptive traits. Then, using the planting site as the focal point, they produced a unique seed zone for each planting site. Although the concept is the same, seed transfer guidelines are often referred to as focal point systems today (Johnson et al. 2004).

In the focal point system, the deployment area is generally defined by seed transfer guidelines based on “geographic, adaptive, ecological, or climatic” criteria (Ukrainetz et al. 2011). However, the main difference between earlier seed transfer guidelines and the focal point approach is the increased use of climate variables (Parker and van Niejenhuis 1996, Ukrainetz et al. 2011), which was facilitated by the development of climate interpolation models (Daly et al. 2008, Wang et al. 2016a, Fick and Hijmans 2017). Rather than restricting seed deployment to a particular geographic distance or change in elevation, seedlots are planted only at sites that have ‘matching’ climates. Matching climates are those with a climatic distance less than or equal to the climate distance threshold (CDT), which is the maximum distance one can move seed without incurring unacceptable maladaptation. Others refer to the CDT as a ‘critical seed transfer distance’ (Ukrainetz et al. 2011) or ‘transfer limit’ (St.Clair et al. 2022). For finding acceptable planting sites, the origin of the seedlot is the focal point. For finding acceptable seedlots, the planting site is the focal point. In either case, matching climates consist of all contiguous and non-contiguous areas that fall within the CDT.

To combine the operational ease of fixed zones and climate-based deployment of the focal point system, Ukrainetz et al. (2011) suggested a hybrid system called the ‘focal zone’ system. The focal zone system uses many small, fixed zones but allows transfers between zones based on matching climates. Although focal zone systems can be more challenging to use than fixed zone systems, they generally lead to larger deployment areas and facilitate climate-based seed deployment and assisted migration. The British Columbia (BC) Ministry of Forests led a formal stakeholder evaluation of the fixed zone, focal point, and focal zone systems for use in BC (O’Neill et al. 2017). Based on seven key objectives, the focal zone approach was ranked as the best alternative, meeting “all objectives well, scoring somewhat better than the focal point system, and much better than the fixed zone system” (O’Neill et al. 2017). Based on similar feedback from members of the Pacific Northwest Tree Improvement Research Cooperative (PNWTIRC), we designed a focal zone deployment system for the Pacific Northwest (PNW).

Climate-based seed deployment is becoming increasingly important because it facilitates assisted migration, the intentional movement of species or populations to areas where they are expected to do well in future climates (Pedlar et al. 2012, Williams and Dumroese 2013). Because of rapid climate change, tree populations are predicted to become progressively maladapted to their local climates (Wang et al. 2006, St.Clair and Howe 2007, Frank et al. 2017), resulting in declines in growth, changes in suitable habitat and species composition, and increases in pests (Iverson and Prasad 2002, Wang et al. 2016b, Wilhelmi et al. 2017). By 2020, global surface temperatures had already increased by about 1.09°C compared to pre-industrial times, and warming will likely or very likely exceed 1.5°C by 2040 (Lee et al. 2023). By 2100, increases in global temperatures are projected to range from 1.4°C to 2.7°C, accompanied by increases in the frequency and intensity of climatic extremes (Lee et al. 2023). Thus, assisted migration via species range expansion or within-species seed transfer will be necessary to achieve well-adapted populations of forest trees (Rehfeldt et al. 2001, Chmura et al. 2011, Nathan et al. 2011, Schreiber et al. 2013). Assisted migration can be integrated into existing reforestation practices by using climate-based deployment of native seed sources, orchard seedlots, or other reforestation materials (Williams and Dumroese 2013).

Two questions must be answered before one can implement climate-based seed transfer. The first question is, ‘Which variables should be used to measure climate distance’? A common approach is to infer important climate variables from provenance tests established in the nursery or field (e.g., St.Clair and Howe 2007, Frank et al. 2017, Prevéy et al. 2018, St.Clair et al. 2019). Although trends are clear, results often vary because of differences in the climates sampled (e.g., provenances or test sites) or variables analyzed. After establishing a distance function using one or more climate variables, the second question is, ‘How far can seedlots be moved?’ The best approach is to infer CDTs from long-term provenance tests using a range of analytical approaches that have been developed (Raymond and Lindgren 1990, O’Neill et al. 2014, Chakraborty et al. 2015). However, the provenance tests needed to infer CDTs directly from responses in the field are limited in number and geographic extent. When genetic tests are unavailable, climate variation within existing deployment zones can be used to infer minimum safe CDTs. For example, because fixed zone systems have generally controlled maladaptation and maintained productivity over many decades, we assume that climate variation within these zones does not exceed acceptable CDTs.

Our overall goal was to develop a climate-based focal zone system to guide seed deployment and assisted migration in the PNW. Key objectives were to facilitate seed transfers among states, provinces, and organizations that are now using different zone sets, and to develop a web application called Zone Matcher that can be used to evaluate deployment options. To accomplish this, we (1) identified zone sets of interest to stakeholders, (2) developed a multivariate climate distance function for guiding tree deployment, (3) inferred safe transfer distances from analyses of provenance tests and climate variation within existing fixed zones, (4) evaluated the performance of our zone system with and without assisted migration, and (5) published the Zone Matcher web application.

## METHODS AND MATERIALS

### Geographic region and zone sets

Our analyses focused on zone sets in BC, California, Idaho, Oregon, Washington, western Montana, and western Wyoming, which we refer to as the PNW (Figure 1). Geographic zone sets are subdivided into zone units, which are contiguous geographic areas. These zone units are subdivided into non-contiguous zones based on elevation. We also studied ecological zone sets that consist of non-contiguous ecological zones not organized into zone units (Table 1). Zone sets were either species-specific or generic, meaning that any tree species can be considered for transfer among these generic zones.

**Figure 1.**
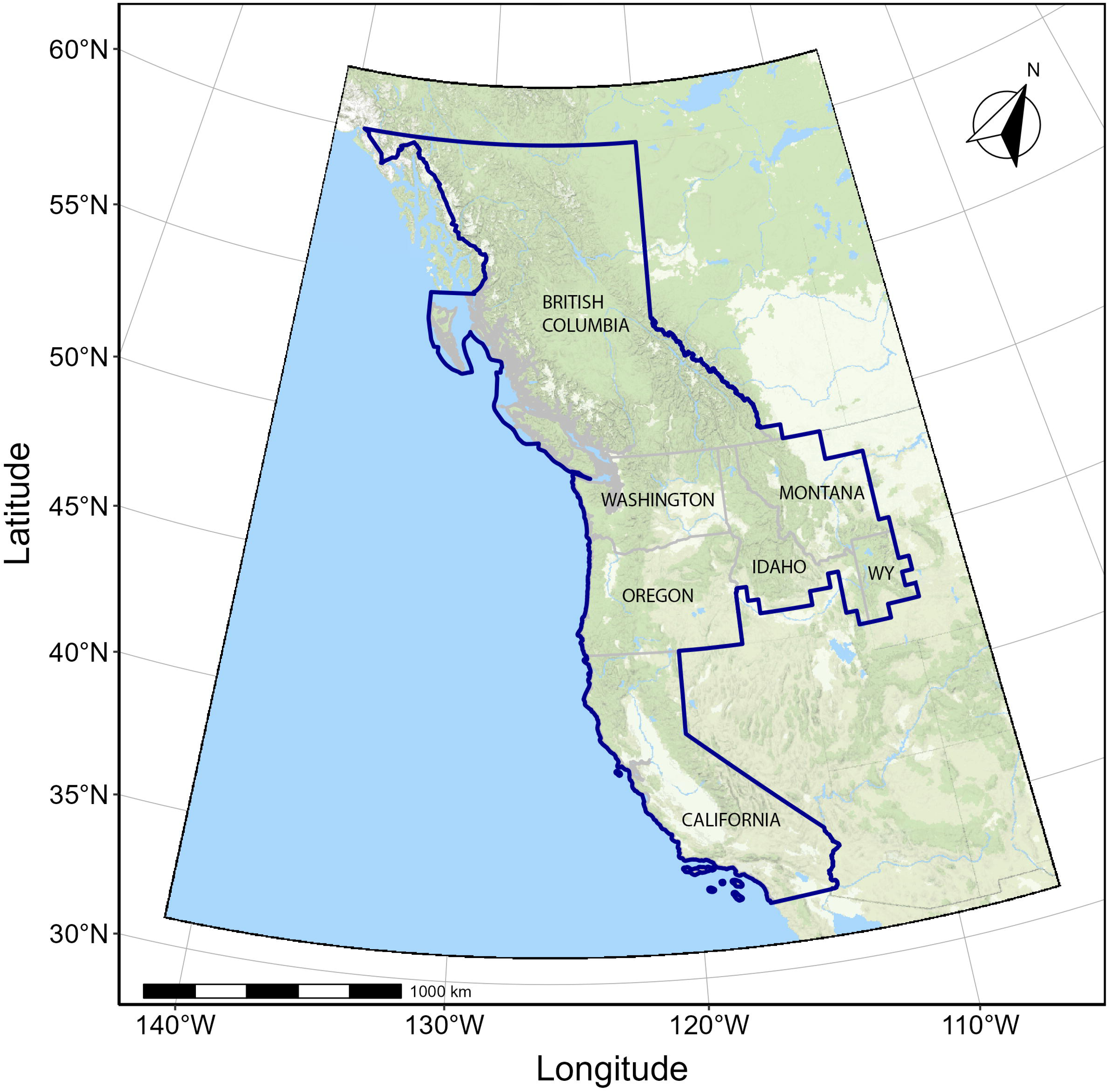
Pacific Northwest study area. We analyzed overlapping zone sets for the Pacific Northwest region of North America. These zone sets covered British Columbia, Washington, Oregon, California, and parts of Idaho, western Montana, and western Wyoming (WY).

For detailed analyses, we selected 6 generic and 10 species-specific zone sets in collaboration with members of the PNWTIRC (Table 1). We obtained existing shapefiles for all zone sets except for Idaho and western Montana (ID/MT). We created the ID/MT zone set using the digital elevation model (DEM) described below to generate 0.5-degree zone units for forested areas in Idaho and western Montana. These zone units were then divided into elevation bands of 500 feet (152.4 meters). We rasterized the shape files and obtained climate data using a DEM from which large bodies of water were excluded. We constructed the DEM using data from the USGS GMTED2010 Viewer (https://topotools.cr.usgs.gov/GMTED_viewer/viewer.htm) (U.S. Geological Survey 2010). The resulting raster had a resolution of 15 arc seconds using the WGS84 geographic coordinate system, which is equivalent to a resolution of ∼450 m, depending on location. At this resolution, the geographic area represented by a single pixel averaged ∼14.1 ha (∼34.9 acres). To obtain rasters for the zone sets, we reprojected the polygon for each zone unit to match the DEM, and then rasterized the polygons using the Geospatial Data Abstraction Library (GDAL; https://gdal.org/) with Bresenham’s line algorithm (Bresenham 1965). This algorithm included some pixels that overlapped with the zone unit boundary, and because we rasterized each zone unit separately, some pixels were claimed by multiple zone units. All geographic zone sets except ID/MT had pre-defined elevation bands which we used to divide the zone units into zones. Then, if these zone units had areas above or below the pre-defined elevations, we divided these into 500-foot increments. The ecological zone sets had no zone units, and their non-contiguous zones were based on either biogeoclimatic ecosystem classification (BEC) or a composite of ecological features (EPA4), rather than on elevation (Omernik and Gallant 1986, Meidinger and Pojar 1991). The BEC system consists of a hierarchy of classification levels (zones, subzones, and subzone variants) based on climate, plant community composition, and soil characteristics (Meidinger and Pojar 1991). We refer to the subzone variants as ‘zones’ to be consistent with the terminology used for the geographic zone sets. Using the raster data described above, we calculated the areas of pixels, zones, and zone units for each zone set using the cellSize function of the *terra* R package (Hijmans 2024).

### ClimateNA data

We used data from ClimateNA v5.60 to develop the climate distance function that was used to identify zones with similar climates (Wang et al. 2016a; http://climatena.ca). After developing the climate distance function, we used data from ClimateNA v7.42 for subsequent analyses. For the climate distance function, we used the DEM described above to interpolate ‘recent climate’ data (1981-2010) for 16 annual climate variables (Appendix S1: Table S2). The DEM was clipped to the PNW study area, converted to ASCII raster format, and then used as input to ClimateNA v5.60 to obtain climate predictions for 1981-2010. After designing the climate distance function, we generated new data using ClimateNA v7.42 for 6 historical periods (1941-1970, 1951-1980, 1961-1990, 1971-2000, 1981-2010, and 1991-2020) and 12 projected period-scenario combinations. These combinations consisted of 3 time periods (2011-2040, 2041-2070, and 2071-2100) and 4 Shared Socioeconomic Pathways (SSP1-2.6, SSP2-4.5, SSP3-7.0, and SSP5-8.5). For the SSPs, the numbers after the dashes are projections of climate forcing in W/m^2^. Climate projections were based on an ensemble of eight global climate models from the Sixth Coupled Model Intercomparison Project (CMIP6; Mahony et al. 2022).

### Zone climate statistics and zone climate samples

In the next eight sections (i.e., up to and including ***Climate distance thresholds (CDTs)***), we describe analyses used to design the 9-variable climate distance function. We conducted these analyses using the 1981-2010 climate data from ClimateNA v5.60. First, we used Python scripts and the numpy.percentile function to extract climate data for each pixel and calculate climate statistics for each zone. For each climate variable and zone, we calculated the difference between the 1st and 99th percentiles (RANGE) and the zone center value (CENTER), which is the mid-point between the 1st and 99th percentiles. We used a similar approach to calculate the center point latitude (LAT) and longitude (LONG) for each zone unit (or zone for ecological zone sets). For the geographic zone sets, we used the LAT and LONG CENTER of the zone unit as the geographic center of the corresponding zones. We then summarized the geographic characteristics of each zone set, including the number of zones, zone areas, and elevation band widths (i.e., zone elevational ranges). We used zone samples for some of the analyses described below. To obtain the zone samples, we sampled zones randomly without replacement by extracting climate data for all pixel locations within a given zone, randomly shuffling them, and then taking the first 1,000 entries from the shuffled dataset.

### Identification of non-forest zones

We removed non-forest zones for some analyses. Non-forest zones were identified using LANDFIRE data and information from the BC Ministry of Forests. For the U.S. zone sets, we used LANDFIRE Remap Canopy Height (CH) data (https://www.landfire.gov/ch.php; Earth Resources Observation and Science Center (EROS) and U.S. Geological Survey 2020). Within the LANDFIRE Remap Existing Vegetation Height (EVH) dataset, pixels with a CH < 1.8 meters are considered non-forest (U.S. Department of Agriculture Forest Service and U.S. Department of the Interior). We aggregated and projected the LANDFIRE raster data (resolution ∼30 m) to match the climate and zone raster data using the *terra* R package, and then flagged zones as non-forest if they had less than 10% of the zone area classified as forest. For the BEC zone set, we removed zones that were described as lacking trees (Meidinger and Pojar 1991), which consisted of zones falling within the four BEC alpine zones: Alpine Tundra (AT), Boreal Altai Fescue Alpine (BAFA), Coastal Mountain-heather Alpine (CMA), and Interior Mountain-heather Alpine (IMA) (British Columbia Ministry of Forests and Range 2006, Forest Service British Columbia Research Branch 2007).

### Filtered and normalized climate data

We removed zones that we identified as outliers based on the RANGE variables (McClure 2021) and removed non-forest zones for some analyses. For the non-normalized data, we identified and removed 176 outlier zones (3.1%) among the six ecological and geographic zone sets (McClure 2021). The climate raster data were then normalized across the entire PNW study area using the yeojohnson function from the *bestNormalize* R package. We did not remove outliers from the analyses of normalized data. Where stated, we removed non-forest zones for analysis; otherwise, non-forest zones were included.

### Correlations among climate variables

Using the forested zones and normalized data, we calculated correlations among the CENTER values for climate variables, LAT, LONG, and photoperiod (PHOTO). We calculated PHOTO for each location (pixel) based on the date mid-way between the summer solstice and autumnal equinox using the photoperiod function from the *meteor* R package. We used the four non-overlapping geographic zone sets (CA, ID/MT, OR66, and WA66; Table 1) and the corr.test function from the *psych* R package to calculate climate correlations and *p-*values.

### Hierarchical partitioning of climate variation

For each of the four non-overlapping geographic zone sets, we partitioned the climate variance among zone units, among zones within zone units, and within zones. For these analyses, we used normalized climate data from a random sample of 1,000 points from each forested zone, and then estimated variance components at each hierarchical level using the lmer function (REML = T; optimizer = “bobyqa”) from the *lme4* R package. After partitioning the variance for each zone set, we averaged the variances (proportions of total) across the four zone sets.

### Random forest analysis

Using the normalized CENTER data from ClimateNA v5.60, we performed random forest classification and regression analyses on the four non-overlapping geographic zone sets. The purpose was to identify climate variables associated with regional climate variation among zone units (classification) and local variation among elevations (regression). Non-forest zones were excluded for this analysis.

First, we used classification analysis to identify which variables were strongly associated with climate variation among zone units. Specifically, we developed a random forest classification model using the *randomForest* R package to predict the zone unit based on a given zone’s climate, with *ntrees* = 1000 and default *mtry* value = 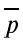 where p is the number of variables (Breiman 2001, Liaw and Wiener 2002). Variables with large importance scores were preferentially included in our climate distance function. For each analysis, the classes were the sampled zone units (N = 20), observations were the zones within zone units, and the predictors were the 16 normalized climate CENTER variables (Appendix 1: Table S2). We randomly sampled 5 zone units from each of the 4 zone sets for a total of 20 zone units per analysis. This analysis was repeated 100 times by sampling new zone units with replacement, resulting in a total of 2,000 zone units analyzed for each replication. We used the out-of-bag (OOB) classification error to choose the final number of model variables, and then ranked the climate variables using the mean decrease in accuracy (MDA) as the variable importance score. We started with all 16 climate variables, and then repeated this analysis 16 times, removing the climate variable with the lowest MDA importance score each time. After replicating this entire process six times, we averaged the MDA importance scores and OOB error for the models containing eight climate variables. We focused on these models because models with eight or fewer variables consistently had near optimal values for OOB error.

In the regression analysis, we used the randomForest function to identify which climate CENTER variables were most closely associated with elevation (default mtry = p/3; Liaw and Wiener 2002). We used a sampling scheme like the one described above, except we used a sliding window approach to sample five adjacent zone units along a latitudinal gradient (i.e., we ranked zone units from southernmost to northernmost). We also used the same approach to sample along a longitudinal gradient (i.e., we ranked zone units from westernmost to easternmost). For each new sample, four of the five zone units overlapped the previous sample. After each run, we added one and subtracted one adjacent zone unit until we had sampled all zone units in the four non-overlapping geographic zone sets. As described above, we averaged variable importance scores across all samples. For the regression analyses, these importance scores consisted of the percent Increase in Mean Square Error (*%IncMSE*) and Increase in Node Purity (*IncNodeP*). As with the classification analysis, we started with 16 climate variables, and then repeated the same analysis after successively dropping the variable with the lowest average *%IncMSE*. After repeating this process six times, we averaged *%IncMSE*, *IncNodeP*, and *pseudo R^2^* (*RSQ*) for the models containing eight climate variables. We focused on these models because models with eight or fewer variables consistently had near optimal values for *RSQ*.

### Climate variable selection and climate distance

We selected climate variables for calculating climate distance based on the correlation analysis, hierarchical partitioning of climate variation, random forest classification, random forest regression, and knowledge of predictive performance of climate variables from genecological studies. We focused on random forest results from the non-overlapping geographic zone sets because classification errors were lower and the *RSQ* values were higher compared to the ecological zone sets. Within the geographic zone sets, we focused on the latitude-based regressions because these had higher *RSQ* values (0.96) compared to the longitude-based analyses (0.87).

For both the classification and regression analyses, models with eight climate variables were optimal or near optimal, so we used these as a starting point for selecting climate variables. However, the classification analyses identified nine unique climate variables among the six random forest replications. Of these, we chose the top six to include in our final combined model (TD, MSP, MAP, EMT, FFP, MCMT). The regression analysis identified eight unique climate variables among the six replications. From these, we chose the top five (MAT, DD_0, DD5, EREF, MCMT), resulting in a combined set of ten unique variables. Finally, we used the correlation analyses, variance partitioning results, and additional random forest analyses to prune the variables to a smaller set of nine. We removed DD_0 because of its strong correlation with MCMT (r = –0.99) and replaced DD5 with MWMT (r = 0.96). The final set of nine variables (EMT, EREF, FFP, MAP, MAT, MCMT, MSP, MWMT, and TD) gave nearly optimal performance for the classification analysis (OOB = 0.16) and regression analysis (RSQ = 0.96) using the same analytical approach as described above.

We assessed whether the nine final climate variables should be weighted differently from one another in the final measure of climate distance by calculating scaled variable importance weights for each climate variable using *MDA* / Σ*MDA* for the classification analysis and *%IncMSE* / Σ*%IncMSE* for the regression analysis. We calculated a combined weight as the mean of its classification and regression weights. Because the nine climate variables had similar combined weights, they were weighted equally for calculating climate distances. We used ClimateNA v7.42 values for these nine climate variables to calculate the Euclidean distances used in subsequent analyses.

### Climate distance thresholds (CDTs)

We used the mid-elevation forested zones to infer CDTs. Preliminary analyses indicated that mid-elevation zones had larger areas and larger RANGE values for MAT and MAP. Thus, for each zone unit, we ranked the zones by elevation and then used the middle half (i.e., interquartile range of zone ranks) to calculate median CDTs for each zone set. We calculated CDTs for the individual climate variables using the non-normalized (absolute) and normalized data (ClimateNA v7.42, 1981-2010), resulting in two CDTs (CDT_ABS_ and CDT, respectively) for each climate variable. For these analyses, we normalized the data using the std_index function of the *SEI* R package (Allen and Otero 2023), and CDTs were calculated as RANGE/2, where RANGE is the difference between the 99^th^ and 1^st^ percentiles for the zone. We also calculated CDTs for four multivariate sets of variables. Sets of 2, 4, 9, or 16 climate variables were selected using the variable importance scores from the random forest analyses. These CDTs (CDT_2_ to CDT_16_) were calculated using normalized climate variables from ClimateNA v7.42 (1981-2010). Multivariate CDTs were defined in the same way as for the individual climate variables (i.e., RANGE/2), but we had to approximate RANGE (= RANGE*) because some zones were too large to calculate percentiles directly. For example, one BEC zone had 863,709 pixels, which would have required calculating more than 3.0 trillion distances for this zone alone (i.e., across all four multivariate sets of variables). Thus, we calculated RANGE* using the following approach. First, to approximate percentile filtering, we removed 2% of points that were farthest from the zone marginal median (i.e., vector of univariate medians). After filtering, we estimated the largest within-zone distance using two methods. For zones with less than 20,000 pixels, we calculated the largest distance among all pairs of points directly using the R dist function. For zones with more than 20,000 pixels, we identified extreme points using the convhulln function from the *geometry* package in R. Because it was impractical to find the convex hull for more than three climate variables simultaneously, we cycled through all climate variables three at a time and combined their convex hull points. In various trials with up to 50,000 points, the maximum distance among all points was nearly identical to the maximum distance calculated using this subsample. To this set of points, we added a subsample of 20,000 points that were farthest from the zone centroid, and then calculated RANGE* as the maximum distance among the combined set of extreme points.

We calculated multivariate CDTs for each zone as RANGE*/2, and then calculated median CDTs for each zone set. Finally, we averaged the median CDTs for the two ecological zone sets and the four geographic zone sets. This was done for the individual climate variables (i.e., CDT_ABS_ and CDT) and multivariate climate sets (i.e., CDT_2_ to CDT_16_).

Finally, we compared our CDTs to the ‘critical seed transfer distances’ (CSTDs) used by the BC Ministry of Forests. We used CSTDs derived from provenance tests of eight coniferous tree species in BC: grand fir (*Abies grandis* (Dougl. ex D. Don.) Lindley), Engelmann spruce (*Picea engelmannii* Parry ex Engelm.), lodgepole pine (*Pinus contorta* Dougl. ex Loud.), western white pine (*Pinus monticola* Dougl. ex D. Don.), ponderosa pine (*Pinus ponderosa* Lawson & C. Lawson), Douglas-fir, western redcedar (*Thuja plicata* Donn ex D. Don.), and western hemlock (*Tsuga heterophylla* (Raf.) Sarg.). The BC Ministry of Forests calculated CSTDs essentially as described by O’Neill et al. (2017), except using an updated Euclidean distance function consisting of latitude and six standardized ClimateNA variables (DD5^0.25^, MAP^-0.75^, MAT, MCMT, MSP^-0.75^, and TD^0.5^) (Greg O’Neill, personal observations). Because the BC CSTDs and our CDTs were measured using different climate distance functions, we developed a regression equation to convert one climate distance to the other. Using raster climate data from the province of BC, we developed and applied a linear regression equation to predict CDTs from the BC CSTDs (*R*^2^ = 0.986) for the eight species listed above.

### Evaluation of climate-based seed deployment

We evaluated climate-based seed deployment using various CD functions, CDTs, and scenarios with and without assisted migration (AM). For these analyses, we used the six partially overlapping zone sets (BEC, EPA4, CA, ID/MT, OR66, and WA66), zone medians of the normalized climate variables, and excluded non-forest zones. Overall, we studied seed deployment across 4,393 partially overlapping zones across a total area of about 252 M ha. The unique area covered by these zones was about 167 M ha.

First, we used the 1991-2020 climate data to compare 5 climate distance functions consisting of 1, 2, 4, 9, or 16 climate variables. For the individual climate variables, we calculated inter-zone CDs (CD_1_) using the differences between zone medians. For the multivariate CDs, we used the Euclidean distances for sets of 2 to 16 variables. Together, the CDTs for the normalized climate variables are referred to as CDT_1_ to CDT_16_. These univariate and multivariate CDTs were calculated as described above. Matching zones were those with an inter-zone CD less than or equal to the corresponding CDT. For example, for each OR66 zone, we calculated the number (N) and area of matching target zones (AREA) for each of the six zone sets. Then, we averaged N and AREA across all OR66 zones to infer how an average OR66 seedlot might be deployed within each zone set and across the PNW overall. These analyses were conducted separately for the geographic and ecological zone sets.

Second, we used the 1991-2020 climate data to evaluate the 9-variable CD function. For these analyses, we used 10 CDTs centered around benchmark CDTs, which we defined as conservative (0.7), moderate (0.9), and liberal (1.1). For example, using the moderate CDT of 0.9, we studied 10 CDTs ranging from 0.225 (= 0.25 × 0.9) to 2.25 (= 2.5 × 0.9), and then used the same multipliers for the conservative (0.7) and liberal (1.1) CDTs.

Third, we used the 9-variable CD function and a moderate CDT of 0.9 to study deployment with and without assisted migration (AM). For the current climate scenario (i.e., ignoring AM), we used the 1991-2020 climate for both the focal and target zones. In the first AM scenario, we assumed the seedlot was adapted to the 1951 to 1980 climate, and the goal was to find matching planting sites using the 2041-2070 climate and a moderate climate change scenario (SSP3-7.0). In the second AM scenario, we assumed the goal was to find matching seedlots for a planting site, using the same seedlot and planting site climates.

Finally, we sampled pairs of zones across all six zone sets, and then calculated climate distances for points sampled from each zone. In total, we sampled 51,850 zone pairs including selfs, and then calculated inter-zone distances for 1,000 pairs of points. Next, we calculated the proportion of inter-zone distances that were ≤ to the CDT (PROP_CDT_). Using CDTs of 0.7, 0.9, and 1.1, we plotted PROP_CDT_ versus the Euclidean distances between the zones. Finally, for each CDT, we developed a prediction equation for PROP_CDT_ using the R *GLM* package and a quasibinomial (logit) distribution. We used six climate distance variables as predictors. These consisted of linear and squared terms of the following three variables: inter-zone distance, standard deviation of within-zone distances for zone 1, and standard deviation of within-zone distances for zone 2.

### Modeled species distributions

We modeled current and projected future species distributions, and then identified zones expected to be suitable for each species. Species distribution models (SDMs) were developed for eight tree species (western larch (*Larix occidentalis* Nutt.), Engelmann spruce, Sitka spruce (*Picea sitchensis* (Bong.) Carrière), lodgepole pine, western white pine, ponderosa pine, Douglas-fir, and western hemlock) using presence-absence data and random forest classification implemented in the *randomForest* R package as described by Zhao et al. (2023). Using these SDMs and ClimateNA v7.42, we derived predicted probabilities of species occurrence (0 to 1 scale), which we call ‘climate suitabilities.’ These values are continuous but were converted to binary classes using a threshold of 0.3 to predict the species’ fundamental niche; that is, values > 0.3 identify locations where the climate is suitable for the species, whereas values ≤ 0.3 indicate locations where the projected climate is unsuitable (Zhao et al. 2023). To obtain zone-level data, we determined the proportion of suitable habitat that is projected to occur in each zone for each future climate scenario. We considered a pixel as suitable habitat if it had a value > 0.3. Then, for each zone, we calculated the proportion of the zone that consisted of suitable habitat. Finally, we considered a zone suitable for a particular species if the zone had ≥ 10% or ≥ 25% suitable habitat. The zone-level results (1 = suitable and 0 = unsuitable) are available in the Excel file that users can download from the Zone Matcher web application.

### Zone Matcher web application

To provide land managers with a user-friendly tool for evaluating zone matches, we developed the Zone Matcher v3.0 web application. ZM was built as a Shiny application in R using the R packages shiny v1.9.1 and leaflet v.2.1.1. The web application was deployed to the shinyapps.io cloud server (RStudio 2024), which can be accessed at https://pnw-focal-zones.shinyapps.io/Zone_Matcher/.

## RESULTS

### Geographic characterization of zone sets

Using BEC ecosystem classifications and LANDFIRE data, we identified 1,270 zones (out of 5,716) that were classified as non-forest in the six partially overlapping zone sets (BEC, EPA4, CA, ID/MT, OR66, and WA66). Compared to the forested geographic zones (mean = 24 K ha), the forested ecological zones were about 15 times as large (mean = 365 K ha) (Appendix S1: Table S3). Because we delineated the zone units of the ID/MT zone set based on a grid system, rather than on topographic characteristics, the ID/MT zone set had a larger number of smaller zones compared to the others. The two Douglas-fir species-specific zone sets, which were designed for Oregon and Washington, had fewer, larger zones than did the generic Oregon and Washington zone sets (Appendix S1: Tables S3 and S4).

### Climate characterization of zone sets

To assess multicollinearity among climate variables, we calculated average correlations among the climate CENTER variables using the non-overlapping geographic zone sets (CA, ID/MT, OR66, WA66) (Appendix S1: Figure S1 and Table S5 upper diagonal). In the upper diagonal, eight pairs of climate variables had strong correlations (*r* ≥ |0.95|), including pairs involving DD_0, DD5, EMT, EREF, EXT, MAT, MCMT, and MWMT. We calculated Pearson correlation coefficients between climate variables and latitude, longitude, and photoperiod (Appendix S1: Table S5) and for the ecological zone sets (Appendix S1: Table S6 and Figure S2). Within our study area, latitude and photoperiod were highly correlated (r = 1.00). We also partitioned climate variation among zone units, among zones within zone units, and within zones (Appendix S1: Figures S3-S8). We then averaged these results across the non-overlapping geographic zone sets (CA, ID/MT, OR66, WA66) to help select the final set of climate variables (Figure 2; Appendix S1: Table S7). We excluded the ecological zone sets from these analyses because they did not have separate zones and zone units. Overall, variation among zone units was greatest for TD, EMT, MSP, MAP, and MCMT (54 to 67% of total variation; Appendix S1: Table S7), making these variables suitable for accounting for large-scale differences in climate. In contrast, variation among zones within zone units was greatest for EXT, EREF, MWMT, DD5, and MAT (58 to 68% of total variation; Appendix S1: Table S7). Thus, we considered these variables suitable for accounting for local differences in climate. Once we selected the final variables for the climate distance function, we calculated empirical CDTs for each zone set (see below).

**Figure 2.**
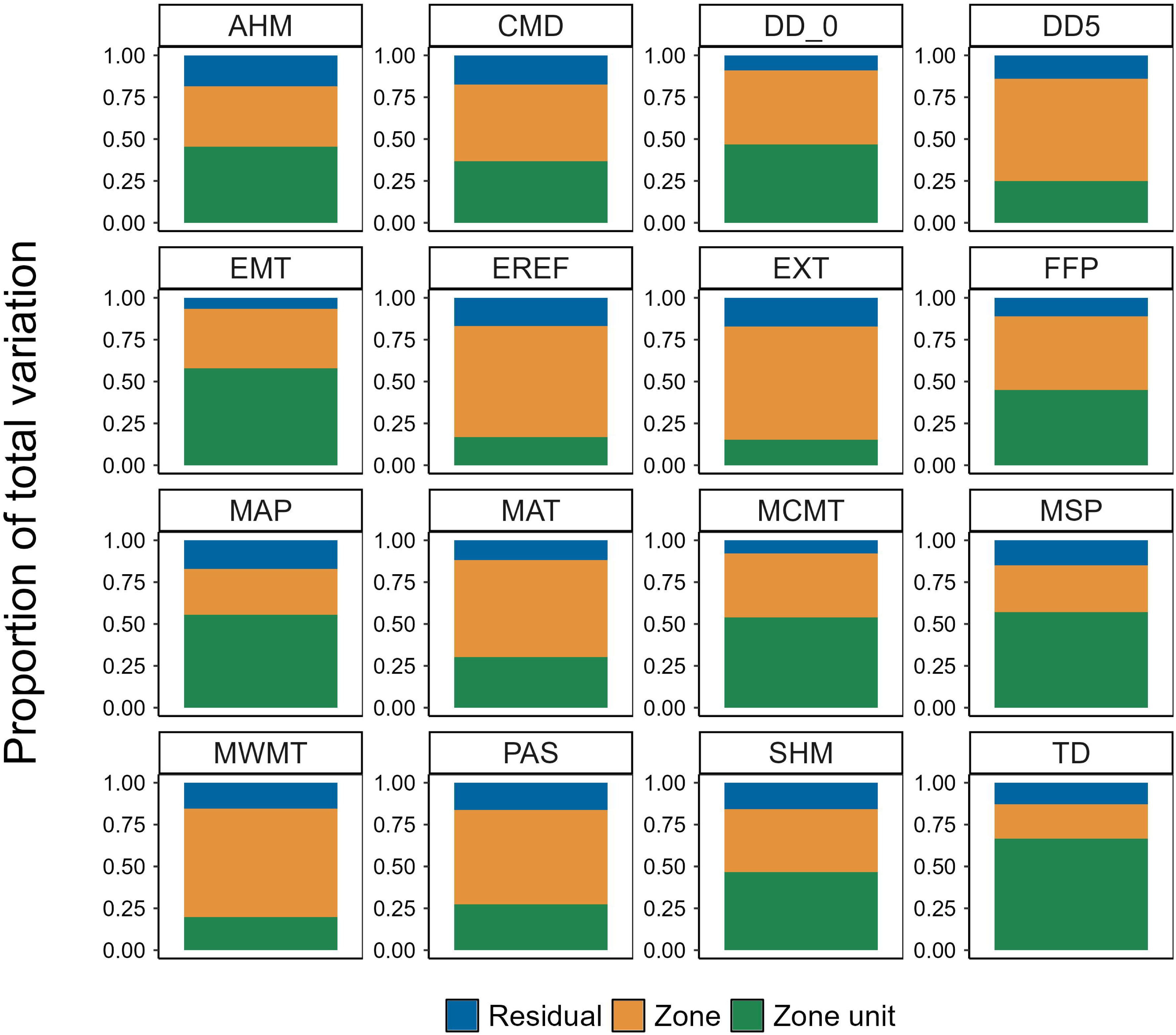
Hierarchical partitioning of climate variation. We partitioned climate variation into three components: within zones (Residual), among zones within a zone unit (Zone), and among zone units (Zone unit), and then expressed the values as proportions of total variation. Figure values are averages across four Pacific Northwest geographic zone sets (CA, ID/MT, OR66, and WA66). Abbreviations for ClimateNA variables are described in Appendix S1: Table S2. Results for individual zone sets are shown in Appendix S1: Figures S3-S8.

### Climate distance

To calculate climate distance, we first selected our final set of climate variables, and then decided how to weight them in a Euclidean distance function. Considering random forest importance scores, climate variable correlations (Appendix S1: Figures S1-S2), and the climate variable partitions (Appendix S1: Table S7), we selected EMT, EREF, FFP, MAP, MAT, MCMT, MSP, MWMT, and TD to calculate climate distance. On average, the random forest models with these nine variables had an OOB error of 0.16 for the classification analysis, and an RSQ of 0.96 for the latitude-based regression analysis (Appendix S1: Table S8). Based on variable importance scores, the most important climate variables were largely reversed between the classification and regression random forest analyses (Table 2). For example, TD was the most important climate variable in the classification analysis but was the eighth most important variable in the regression analysis. Similarly, MAT was the most important climate variable in the regression analysis but was the ninth most important climate variable in the classification analysis. These results suggest that different sets of climate variables are better for characterizing regional variation among zone units (classification analysis) versus local variation associated with elevation (regression analysis). The final nine climate variables included the five variables with the most variation among zone units (TD, EMT, MSP, MAP, and MCMT), and three of the five climate variables with the most variation among zones within zone units (EREF, MWMT, and MAT) (Appendix S1: Table S7).

**Table 2.** Climate variable importance scores and weights from random forest classification (Zone unit) or random forest regression (Elevation).

| Zone unit |  |  | Elevation |  |  | Zone unit + elevation |  |
| --- | --- | --- | --- | --- | --- | --- | --- |
| Clim var* | MDA<sup>†</sup> | Weight<sup>‡</sup> | Clim var* | %IncMSE<sup>§</sup> | Weight<sup>‡</sup> | Combined weights<sup>‡</sup> |  |
| TD | 76.18 | 0.17 | MAT | 22.18 | 0.17 | MSP | 0.13 |
| MSP | 73.49 | 0.17 | EREF | 20.72 | 0.16 | TD | 0.12 |
| MAP | 60.13 | 0.14 | MCMT | 17.33 | 0.13 | MCMT | 0.11 |
| EMT | 55.57 | 0.13 | MWMT | 16.11 | 0.12 | EREF | 0.11 |
| FFP | 51.33 | 0.12 | FFP | 12.81 | 0.10 | EMT | 0.11 |
| MCMT | 42.29 | 0.10 | EMT | 12.72 | 0.10 | MAT | 0.11 |
| EREF | 29.67 | 0.07 | MSP | 11.11 | 0.08 | FFP | 0.11 |
| MWMT | 28.07 | 0.06 | TD | 10.11 | 0.08 | MAP | 0.10 |
| MAT | 22.56 | 0.05 | MAP | 9.63 | 0.07 | MWMT | 0.09 |
| OOB error<sup>¶</sup> = | 0.16 | 1.00 | RSQ<sup>#</sup> = | 0.96 | 1.00 |  | 1.00 |

\*Clim var refers to ClimateNA variables described in Appendix S1: Table S2.
<sup>†</sup>MDA is the mean decrease in accuracy.
<sup>‡</sup>Weights for each climate variable are $MDA / \sum MDA$ and $\%IncMSE / \sum \%IncMSE$ . Combined weights for each climate variable are the mean of zone unit weight and elevation weight.
<sup>§</sup>%IncMSE refers to the increase in mean square error of the predictions when the values for the variable are randomly shuffled.
<sup>¶</sup>OOB error is the out-of-bag error associated with the model.
<sup>#</sup>RSQ is the pseudo R-squared associated with the model ( $1 - MSE/Var(y)$ ).

We used random forest importance scores to judge whether the final set of climate variables should be weighted equally when calculating Euclidean distance. We calculated weights for the classification analysis and regression analysis using the scaled variable importance scores. The combined weights, averaged from the classification and regression weights, varied from 0.09 for MWMT to 0.13 for MSP (Table 2). Thus, because the combined weights were similar, we gave the climate variables equal weight in the Euclidean distance function.

### Climate distance thresholds (CDTs)

For each zone set, we inferred CDT_ABS_ for the individual climate variables (Figure 3; Appendix S1: Table S9) and CDTs for four multivariate sets of variables (Figure 4). Low and high elevation zones tended to be smaller than those at mid-elevations (Figure 5a) and had less climate variation (Figure 5b, c). Thus, to avoid overly conservative estimates of CDTs, we used the mid-elevation zones to calculate CDTs. Generally, the ecological zone sets had larger CDTs than did the geographic zone sets (Figures 3-4; Appendix S1: Table S9). Overall, the median CDT_9_ was 1.10 for the ecological zone sets and 0.66 for the geographic zone sets. Within the ecological zone sets, EPA4 had a larger CDT_9_ than did BEC. Within the geographic zone sets, WA66 had the largest CDT_9_ and OR66 had the smallest.

**Figure 3.**
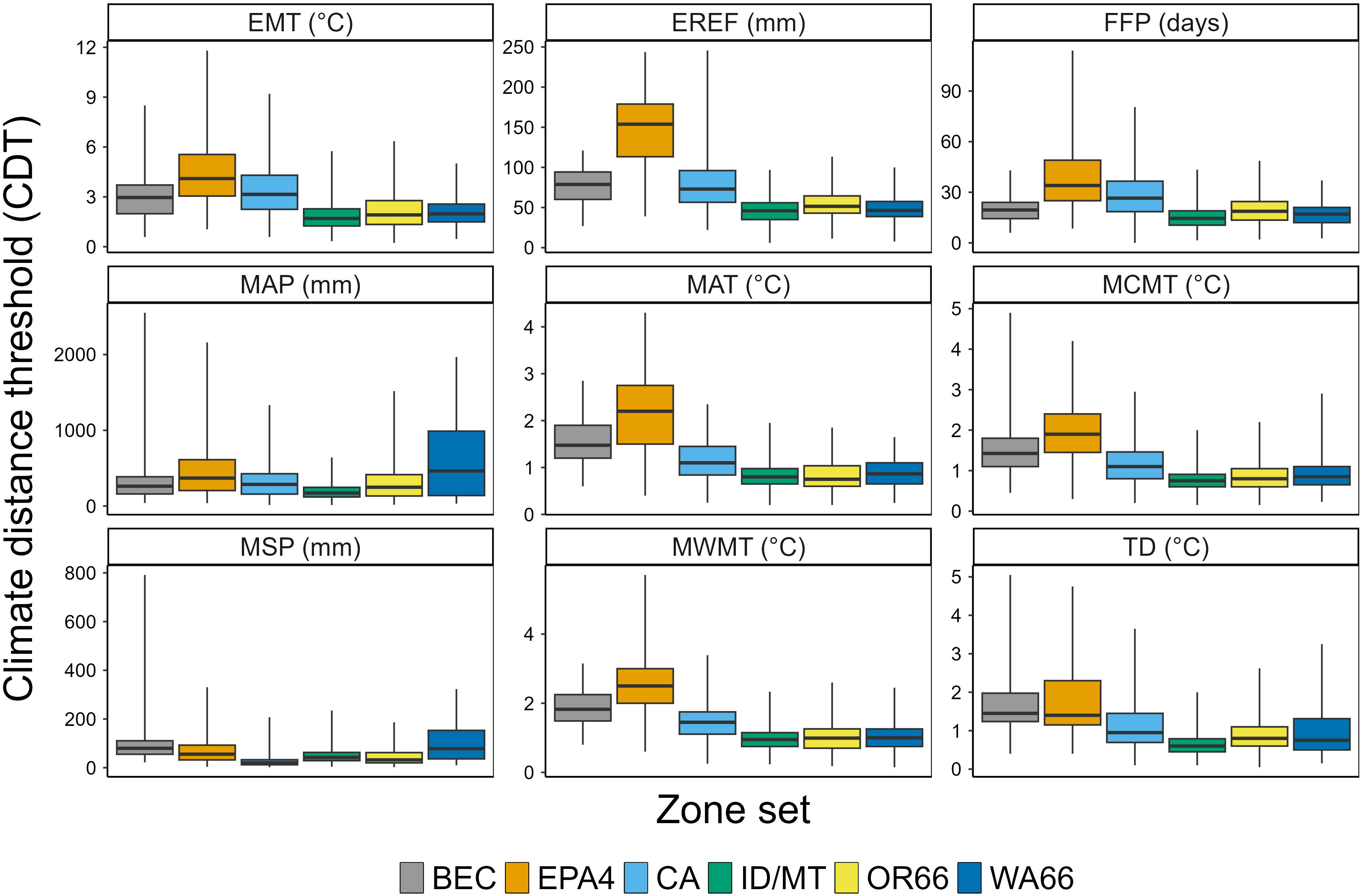
Climate distance thresholds for individual climate variables. We calculated climate distance thresholds (CDTs) for six generic Pacific Northwest zone sets (i.e., non-species-specific zone sets). This figure shows the CDTs for individual climate variables using non-normalized data for the forested mid-elevation zones. CDTs were calculated for each zone as RANGE/2, where RANGE is the difference between the 99^th^ and 1^st^ percentiles. BEC and EPA4 are ecological zone sets, whereas the remainder are geographic zone sets. The black line within each boxplot is the median, the upper hinge of the boxplots is the 75^th^ percentile, the lower hinge is the 25^th^ percentile, the lower whisker shows the minimum value, and the upper whisker shows the maximum value. Abbreviations for ClimateNA variables are described in Appendix S1: Table S2.

**Figure 4.**
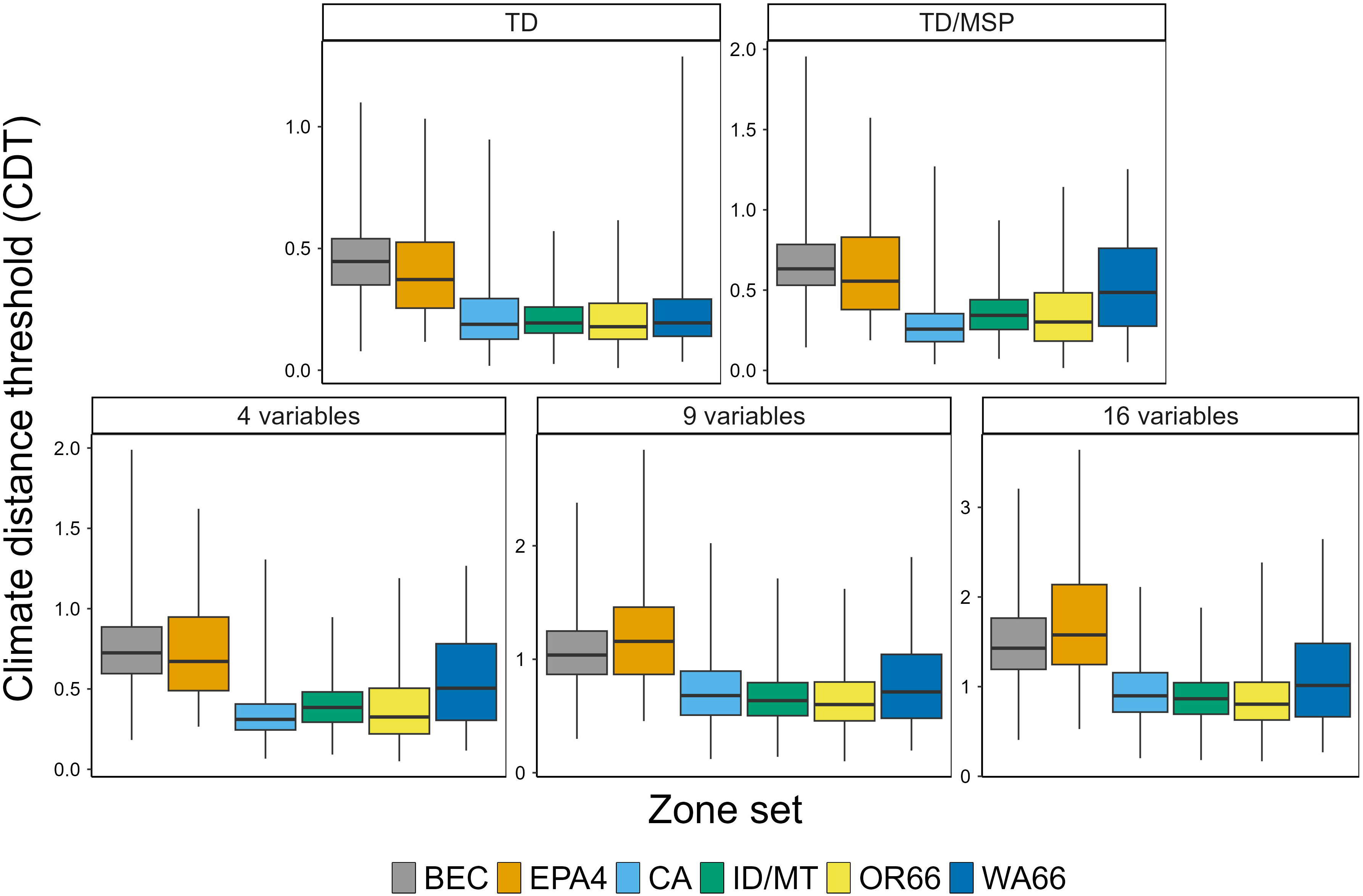
Climate distance thresholds for multivariate climate distance functions. Climate distance thresholds (CDTs) were summarized by zone set for one univariate distance function and four multivariate Euclidean distance functions. The 4-variable function included TD, MSP, MCMT, and EREF, the 9-variable function included our final set of 9 climate variables, and the 16-variable function includes all 16 ClimateNA variables we studied. Abbreviations for ClimateNA variables are described in Appendix S1: Table S2. Boxplot statistics are described in Figure 3. CDTs can be compared among zone sets but not among distance functions because the distance functions have different units.

**Figure 5.**
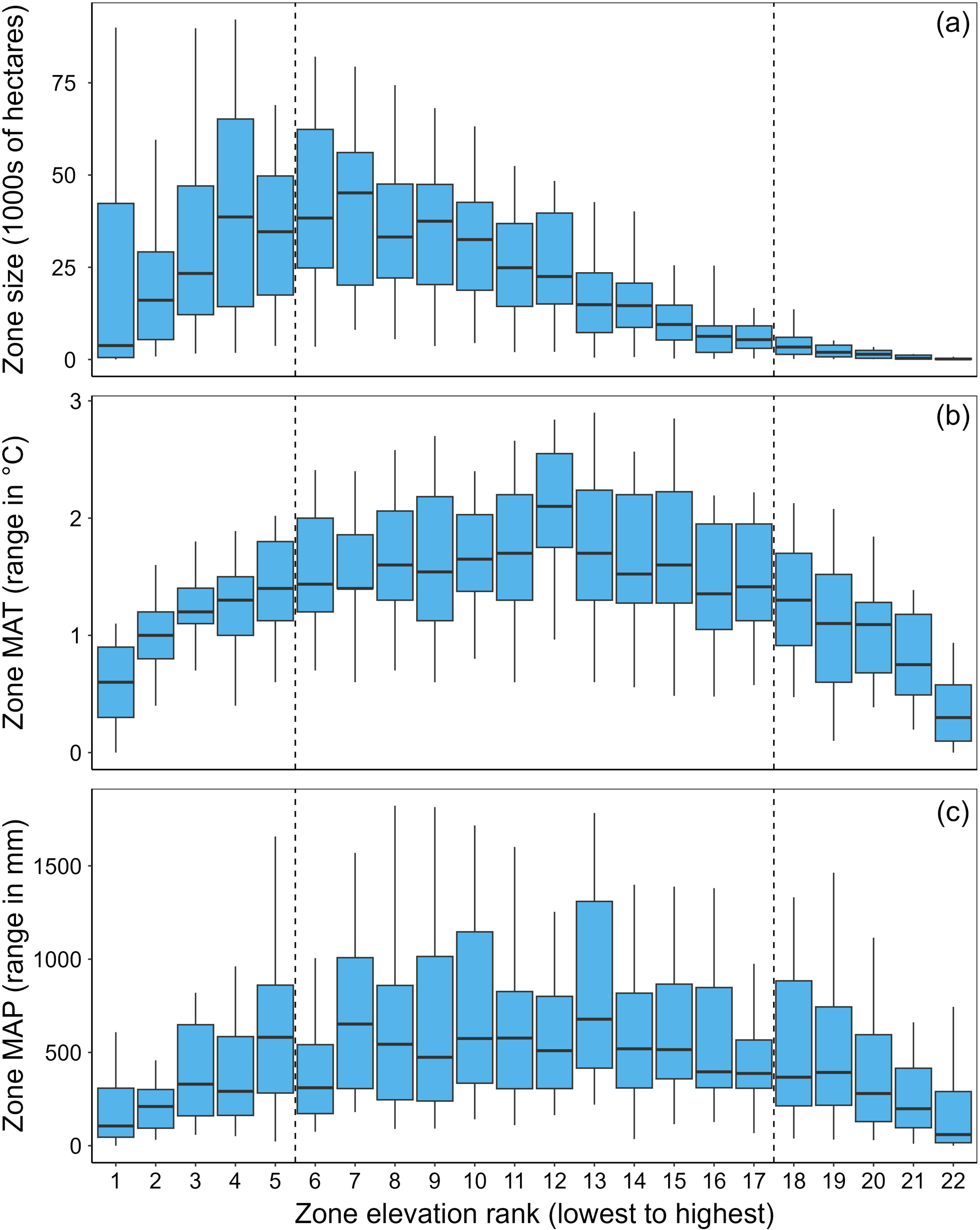
Zone size (a), mean annual temperature (MAT) (b), and mean annual precipitation (MAP) as a function of zone elevation rank for the OR66 zone set. OR66 zones were ranked by mean elevation within zone units (x-axis). The y-axes are zone area, panel (a), within-zone RANGE in MAT, panel (b), and within-zone RANGE in MAP, panel (c). RANGE is the difference between the 1^st^ and 99^th^ percentiles. The black line within each boxplot is the median, the upper hinge of the boxplots is the 75^th^ percentile, the lower hinge is the 25^th^ percentile, the lower whisker shows the minimum value, and the upper whisker shows the 90^th^ percentile. Panel (a) shows that zone sizes are larger for mid-elevation zones. Panels (b) and (c) show that climate variation is greater for mid-elevation zones.

We inferred acceptable CDTs using three approaches. First, we calculated climate variation within existing zones using the nine-variable climate distance function (see CDT_9_ in Appendix S1: Table S9). Second, we studied the relationships between the nine-variable distance function and the non-normalized climate variables (Appendix S1: Figure S9). Finally, we inferred CDTs indirectly from provenance tests by using linear regression to predict CDT_9_ from CSTDs for eight conifer species tested in BC (Appendix S1: Table S10). These CDTs ranged from 0.807 for lodgepole pine to 1.380 for Engelmann spruce. Based on this evaluation, Zone Matcher includes an option for selecting a CDT value of 0.7 (conservative), 0.9 (moderate), or 1.1 (liberal). However, for flexibility, Zone Matcher finds matching target zones using a distance cutoff of 2 × CDT. Nonetheless, the safest approach is to select the closest possible match and only use matches larger than 1.1 when necessary. Finally, the proposed matches should be validated by comparing the climates of the focal and target zones using the individual climate variables as a guide.

### Evaluation of climate-based seed deployment

First, we studied the association between deployment area and number of climate variables in the CD function. For these analyses, we used the 1991-2020 climate data and separate CDTs for the geographic and ecological zones. Although these analyses ignore assisted migration, we report results for scenarios involving assisted migration below. Overall, the deployment area declined as more climate variables were included in the CD function, stabilizing between 4 and 16 climate variables (Figure 6a). For the geographic zones, the deployment area declined from 13.5% of the total area for the single TD variable (CD_1_) to 1.7% for the 9-variable CD function (CD_9_). For the ecological zones, the corresponding values were 25.3% for TD and 5.9% for the 9-variable CD function (Figure 6a).

**Figure 6.**
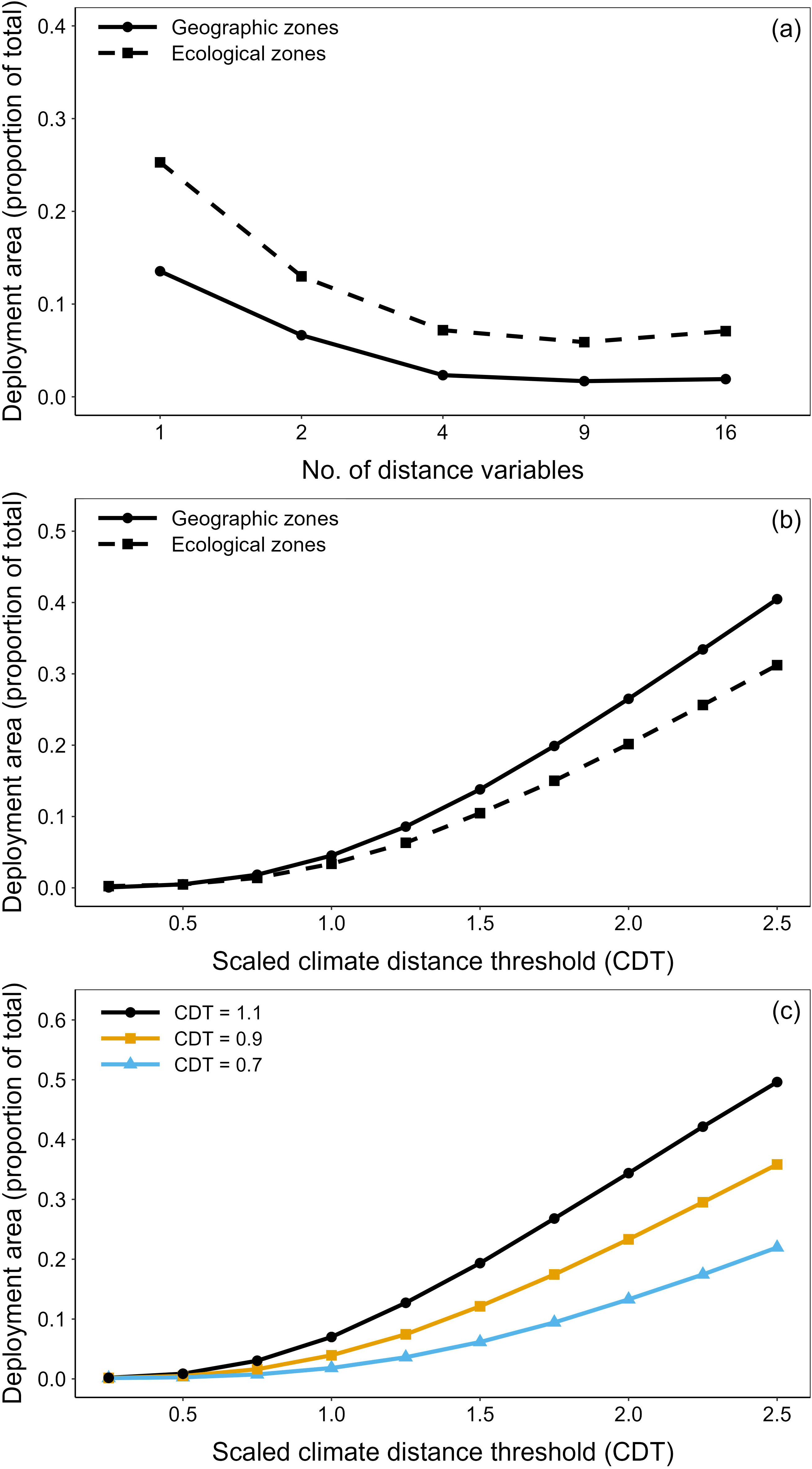
Performance of climate-based focal zones. Panel (a) shows how the mean deployment area as a proportion of the total target area declines with increasing number of climate variables used to build the climate distance function. These analyses were conducted separately for the geographic and ecological zone sets. Total area of geographic zones was 89.2 M ha across 3,940 zones, whereas the total area of ecological zones was 163 M ha across 453 zones. Panel (b) shows matches using the final 9-variable climate distance function and 10 climate distance thresholds (CDTs) ranging from 0.225 to 2.25. For the x-axis, these CDTs were scaled using the ‘moderate’ CDT of 0.9 as a baseline. That is, the x-axis ranges from 0.25 (0.225/0.9) to 2.5 (2.25/0.9). Panel (c) shows matches using the same general approach but using three CDT values (0.7, 0.9, and 1.1) as baselines. In panel (c), the values are the means of the geographic and ecological zone sets.

Next, we used the CD_9_ function to study how the deployment area changed as the CDT was increased from a scaled CD of 0.25 to 2.5 (Figure 6b). The increase in deployment area was nearly linear with CDT, and at a scaled CD of 1.0 (CD = 0.9), the deployment area was 4.52% of the total area for the geographic zones and 3.37% for the ecological zones. Although we used the same CDT for the geographic and ecological zones, the deployment area was consistently larger for the geographic zones. Figure 6c shows the same type of analysis, but using three CDTs (0.7, 0.9, 1.1) based on the options available in the Zone Matcher web application. In contrast to Figures 6a and 6b, Figure 6c shows deployment areas averaged across the geographic and ecological zones.

Table 3 shows the area of matches for focal zones from each zone set using a CDT of 0.9 and scenarios with or without assisted migration. Under ‘Zone set statistics,’ Table 3 shows the total area, number of zones, and the average fixed zone size for each zone set. For example, the ID/MT zone set had the largest number of forested zones (N = 1,807) and smallest average zone size (19.1 K ha). The BEC zone set had the fewest forested zones (N = 200) and largest zone size (407.7 K ha). Table 3 also shows how the focal zone system expands the deployment area compared to fixed zones. If we ignore assisted migration, results are shown in the ‘Current climate’ row. Using the OR66 zones, the deployment area was 23 K ha on average (i.e., the mean fixed zone size), but expanded nearly 100-fold to 2.28 M ha using the focal zone approach. The matching areas were sometimes smaller and sometimes larger using the two assisted migration scenarios (AM1 and AM2; Table 3). In the AM1 scenario, the goal was to find suitable planting sites for a seedlot. In the AM2 scenario, the goal was to find suitable seedlots for a planting site. Although the matching areas were similar for AM1 and AM2, the AM1 area was less than AM2 in the southernmost zone sets (OR and CA). Because matching seedlots are generally associated with climates warmer than the planting site, suitable seedlots are rarer for planting sites near the trailing edge of climate change.

**Table 3.** Performance of the climate-based focal zone system using the 9-variable climate distance function and a climate distance threshold (CDT) of 0.9.

|  | Zone set |  |  |  |  |  |
| --- | --- | --- | --- | --- | --- | --- |
|  | BEC | EPA4 | CA | ID/MT | OR66 | WA66 |
| **(a) Zone set statistics\*** |  |  |  |  |  |  |
| Total area (M ha) | 81.53 | 81.63 | 25.90 | 34.54 | 15.44 | 13.33 |
| Number of zones (N) | 200 | 253 | 900 | 1807 | 678 | 555 |
| Mean zone area (K ha) | 407.7 | 322.6 | 28.8 | 19.1 | 22.8 | 24.0 |
| **(b) Matching area in the PNW (M ha)<sup>†</sup>** |  |  |  |  |  |  |
| Current climate<sup>‡</sup> | 6.81 | 11.36 | 2.01 | 5.62 | 2.28 | 1.93 |
| AM1 - Find plantations<sup>‡</sup> | 5.60 | 4.59 | 2.80 | 5.47 | 4.00 | 4.62 |
| AM2 - Find seedlots<sup>‡</sup> | 5.02 | 4.54 | 2.98 | 5.47 | 4.63 | 4.36 |

\*Total area is the area of all forested zones in the zone set in millions of ha. Mean zone area is the mean zone area in thousands of ha.
<sup>†</sup>Matching area is the unique area of all target zones found for a single focal zone. Matching areas were found for each zone in the zone set, and then averaged.
<sup>‡</sup>The current climate data were derived using the 1991-2020 climate data for the focal and target zones. For the first assisted migration scenario (AM1), we used the 1951-1980 climate as the focal seedlot climate and a projected climate in 2041-2070 as the target plantation climate. For the AM2 scenario, we used the 2041-2070 climate as the focal plantation climate and the 1951-1980 climate as the target seedlot climate. Both projected climates were based on the SSP3-7.0 Shared Socioeconomic Pathway.

Inter-zone transfers are considered acceptable when the distance between zones is less than or equal to the CDT. However, because inter-zone distances are calculated using climate medians, some inter-zone transfers will be less than the CDT and others will be greater. Thus, we calculated the proportion of inter-zone transfers ≤ CDT for a range of inter-zone distances. Using CDTs of 0.7, 0.9, and 1.1, the predicted proportions ranged from about 0.25 to 0.3 (Figure 7).

**Figure 7.**
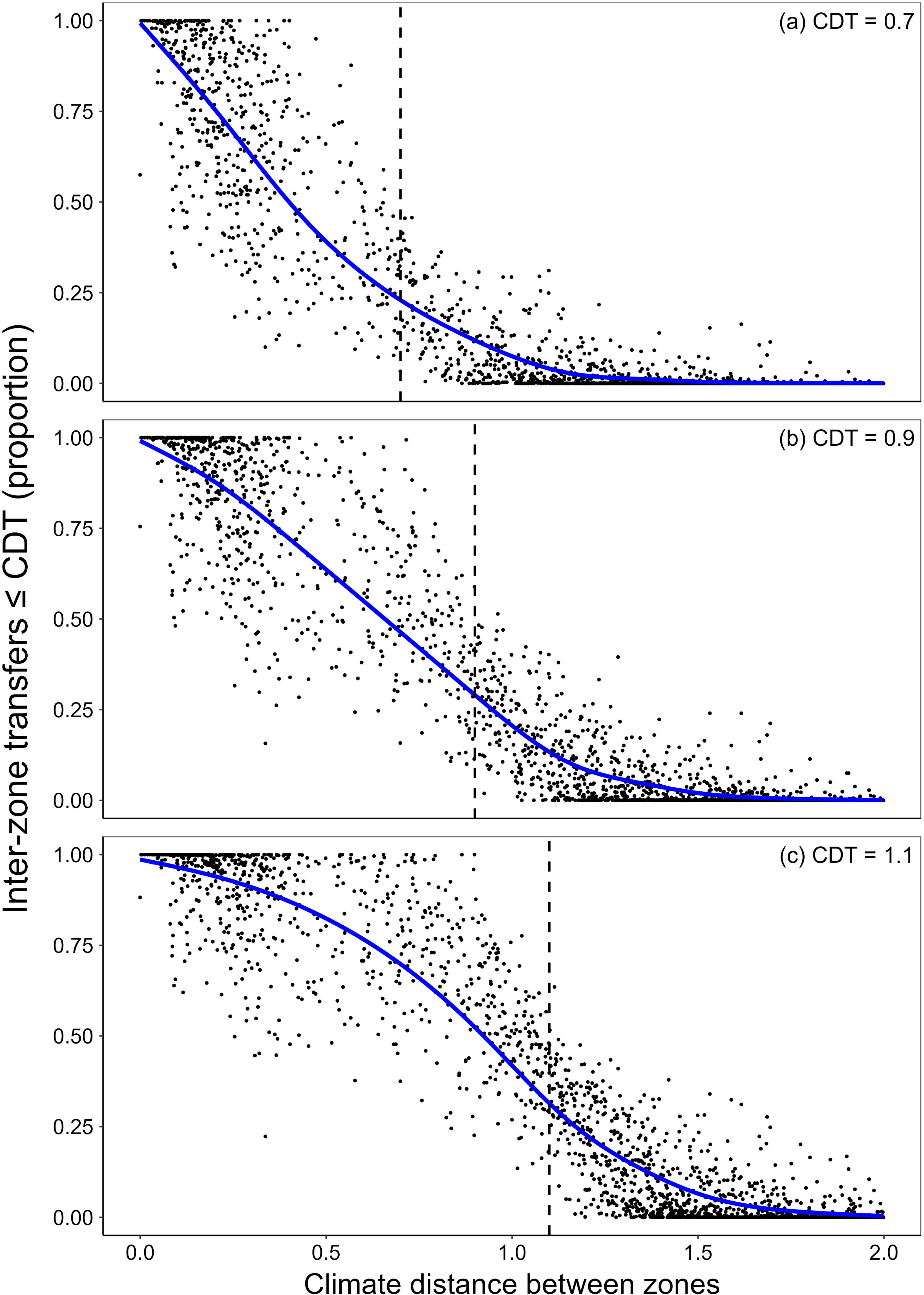
Proportion of inter-zone transfers less than or equal to the climate distance threshold (CDT) versus the climate distance between zones. Results are shown for (a) the conservative CDT = 0.7, (b) moderate CDT = 0.9, and (c) liberal CDT = 1.1. Climate distances between sites in each zone were calculated using a Euclidean distance function comprised of nine normalized climate variables. Distances between zones were calculated using the climate variable medians. The vertical dashed lines show the separate CDTs used for each panel.

### Zone Matcher web application

Using normalized climate variables and the 9-variable climate distance function, Zone Matcher v3.10 calculates climate distances between the selected ‘Focal zone’ and all candidate target zones in the selected zone sets (Figure 8). Matching zones are identified when the CD is less than twice the user-supplied CDT. Users can select a conservative CDT (0.7), moderate CDT (0.9), or liberal CDT (1.1) to generate candidate matches. As a guide, the empirical CDTs shown in Figure 4 are summarized in a table in Zone Matcher, as are the values inferred from the BC CSTDs. The matching zones are displayed on a map and as a list of zones that can be downloaded in an Excel file with other information, including projected zone-level species suitabilities for 8 species and 12 future climate scenarios. Zone Matcher also displays zone median values for 16 non-normalized climate variables, latitude, longitude, and photoperiod. These data are shown for the ‘Focal zone’ and a single selected ‘Comparison zone.’

**Figure 8.**
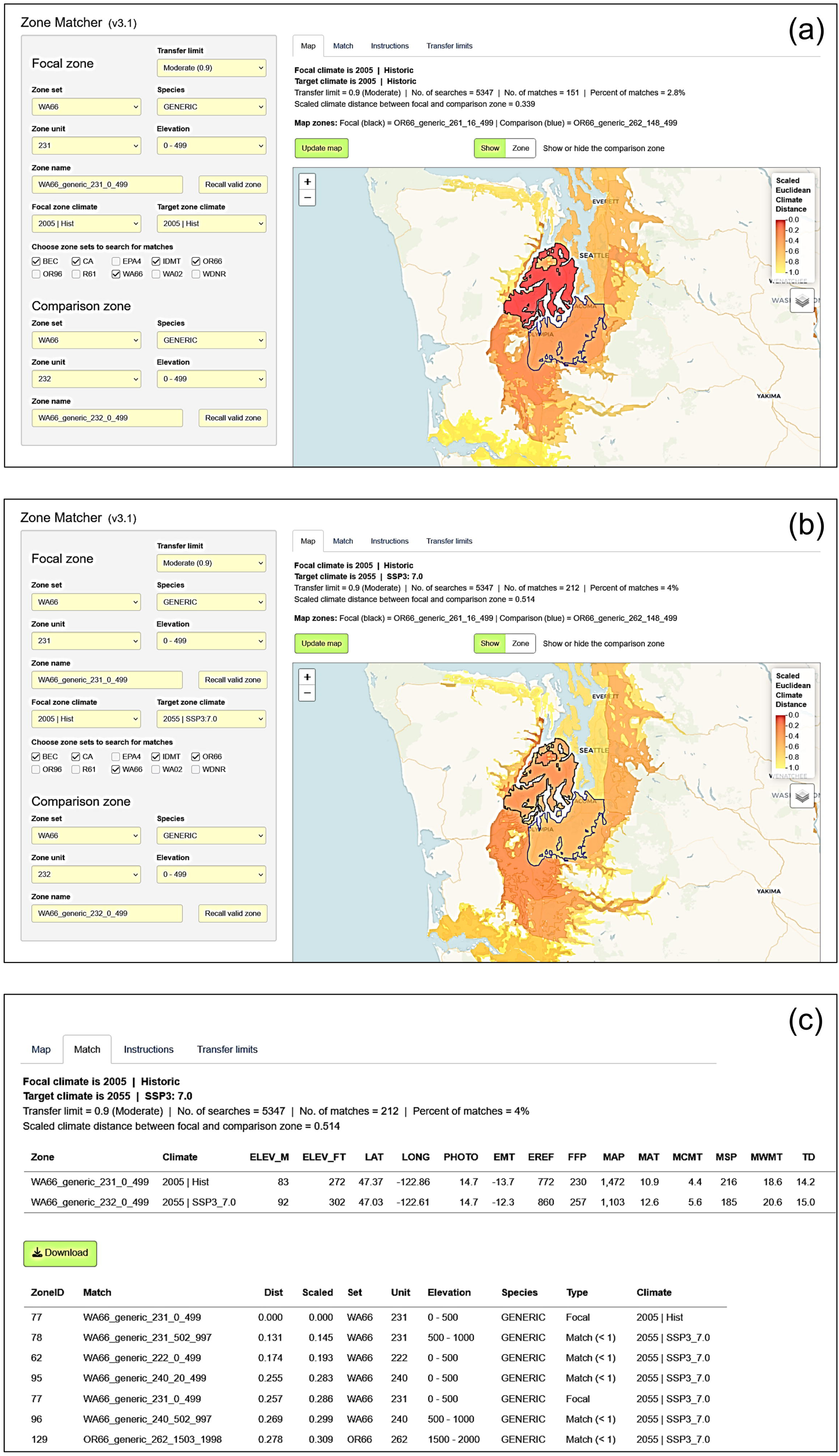
Examples of zone matches identified using the Zone Matcher web application. Matches were found for a WA66 focal zone (black border) in zone unit 231, elevation band 0-499 feet, using the ‘moderate’ climate distance threshold (CDT) of 0.9 Euclidean distance units. Panel (a) shows matches between the 1991-2020 climates (2005 | Hist) of the focal zone and target zones. The target zones include a single comparison zone (blue border), which is WA66 zone unit 232, 0-499 feet. Panel (b) shows matches between the focal zone (2005 | Hist) and the projected climates of the target zones in 2041-2070 using the SSP3-7.0 Shared Socioeconomic pathway (2055 | SSP3_7.0). Panel (c) shows the climates of the focal and comparison zones, and below that, is a list of the top 5 target zones for the listed climates. Matches to the focal zone are displayed from the closest (dark orange) to the farthest (light yellow) climate distance (CD). The “Dist” column shows the absolute CD, whereas the “Scaled” column shows the scaled Euclidean distance (CD/CDT). The Zone Matcher output shown in panel (b) can be downloaded to an Excel file.

## DISCUSSION

We designed a climate-based system for deploying forest tree seed from native populations and seed orchards. Compared to traditional fixed-zone systems, our climate-based, focal zone system substantially expands seed deployment and facilitates assisted migration by incorporating historical climates and future climate scenarios. The system can be implemented using a freely-available web-based application called Zone Matcher. Zone Matcher produces maps of climatically similar zones for three CDT options (conservative = 0.7, moderate = 0.9, and liberal = 1.1), and a list of all matching zones within a scaled CD of 2.0. Because Zone Matcher is a climate matching tool, the Zone Matcher approach and R package code could be adapted to other situations where one wishes to identify and display matching climates using a Geographic Information System (GIS).

### Why use climate-based seed deployment?

One reason to use a climate-based approach is to expand the area of seed deployment compared to fixed zones. Fixed zones cover small to modest-sized areas—only about 24 K ha for the geographic zones and 365 K ha for the ecological zones we studied (Appendix S1: Table S3).

Geographic zones tend to be smaller because they are restricted to elevation bands within contiguous geographic areas (zone units). Ecological zones are usually larger because they include large non-contiguous areas that have similar ecological characteristics (Omernik and Gallant 1986, MacKenzie and Meidinger 2018). Using a focal zone approach and current climate, the deployment area increased about 17-to 35-fold for the ecological zones and 70-to 300-fold for the geographic zones using a CDT of 0.9 (Table 3). Compared to the current climate scenario, the target areas were sometimes substantially different using two assisted migration scenarios (Table 3). Deployment risk can be managed by using a more conservative or liberal CDT, using the CDTs we estimated from different zone sets and provenance tests as a guide. This highlights an important advantage of the focal zone approach—the ability to use a single set of generic zones and then adjust the CDT to account for species differences. In a fixed zone system, species-specific zones may be needed because the zone sizes define the CDTs.

Although the focal zone approach expands seed deployment, the areas suitable for planting may be much smaller than predicted once non-climatic factors are considered—e.g., local factors such as soils, topography, photoperiod, pests, or projected species distributions. Nonetheless, by expanding seed deployment, breeding programs may be able to reduce costs by using fewer breeding populations and fewer seed orchards. For native seedlots, climate-based seed deployment could simplify seed collection, seed inventory, and seedling production. Also, climate-based seed deployment allows more seedlots to be considered for a given planting site. This is particularly important for post-fire revegetation, where limited availability of seed sources is often a problem (Ukrainetz et al. 2011). Because of increasingly large and widespread wildfires in the western U.S., it has become increasingly difficult to find suitable seedlots for reforestation (Fargione et al. 2021).

Climate-based seed deployment also facilitates assisted migration. Because of local adaptation and rapid climate change, native populations of trees will become increasingly maladapted to their local climate (Wang et al. 2006, St.Clair and Howe 2007, Frank et al. 2017). Likewise, breeding populations will become increasingly maladapted to areas where they are currently deployed (Rehfeldt et al. 2001, O’Neill et al. 2017). Thus, to account for shifts in suitable habitat, assisted migration will be needed to maintain forest health and productivity (Rehfeldt et al. 2001, Aitken et al. 2008, Chmura et al. 2011, Nathan et al. 2011, Zhao et al. 2023). Fixed zone systems inhibit assisted migration because seed cannot be transferred among fixed zones. Instead, focal zone deployment systems allow land managers to adjust seed deployment to match future climates. Using the Zone Matcher web application, users can practice assisted migration by matching historic climates with 12 future climate scenarios consisting of 3 future time periods and 4 SSPs.

### Why use a focal zone system?

We developed a focal zone system for climate-based seed deployment because it is simpler to use than a focal point system. Ukrainetz et al. (2011) recommended a hybrid focal zone system with many, non-overlapping fixed zones. When British Columbia developed their climate-based seed deployment system, O’Neill et al. (2017) surveyed stakeholders and similarly found a focal zone system best met criteria for ease of use and ability to be integrated into other decision support tools. In focal point deployment systems, the deployment area differs for every focal point (location), yielding a vast number of zones. Consequently, results are difficult to integrate with other decision support tools. For example, seedlots collected from natural stands are often tracked using the name of the corresponding seed zone. Thus, the list of matching zones in Zone Matcher is also a list of suitable seedlots. Likewise, improved seed from seed orchards may be deployed using the seed zones originally designed for native populations. Because of existing links between seedlots and deployment zones, focal zone systems can be readily adopted by land managers. Focal zone systems also facilitate sharing seedlots among organizations because climate matches can be easily found between zones in different zone sets. For example, the average BC (BEC) zone matched ∼26 zones spanning 6.81 M hectares in Oregon and Washington (Appendix S1: Table S11).

Finally, focal zones facilitate the use of multivariate Euclidean distance functions, which have desirable characteristics (see below). Using our study area and DEM resolution, a focal point system would require calculating at least ∼12.4 M Euclidean distances for a single focal point. Because our focal zone system requires less than 4,140 calculations to cover the same area, including overlapping zones, it is more computationally efficient.

### Developing a climate distance function

We used a multivariate approach to characterize climate distance. The final set of nine climate variables (EMT, EREF, FFP, MAP, MAT, MCMT, MSP, MWMT, and TD) include temperature and precipitation variables, explain small-scale and large-scale differences in climate across the region, and are important indicators of adaptive variation based on genecological studies. To explain large-scale (e.g., regional) variation in climate, we identified variables that accounted for the greatest amount of climate variation among zone units. The top five variables in this category were TD, EMT, MSP, MAP, and MCMT (Appendix S1: Table S7). Furthermore, these variables were among the most important for classifying zones into zone units using random forest classification (Table 2). These results are broadly consistent with known climate variation across the region. TD, MSP, and MAP vary west to east because of the moderating effect of the Pacific Ocean on temperatures (e.g., TD) and the rain shadow effect caused by the mostly north-south mountain ranges (e.g., MSP, MAP). Thus, climate variables that vary from the coastal to interior west are often associated with variation in adaptive traits such as drought hardiness (St.Clair and Howe 2007). Large scale variation was also associated with temperature—in our analyses, we identified extreme minimum temperature (EMT) as one of the more important variables at the regional scale.

Climate variables that explained high levels of variation among zones (i.e., within zone units), EXT, EREF, MWMT, DD5, and MAT, account for smaller-scale climate differences associated with elevation (Appendix S1: Table S7). Random forest regression identified three of these same variables as being important for explaining local differences in elevation (MAT, EREF, MWMT; Table 2). Overall, we identified seven local-scale variables using these two types of analyses, all of which were associated with temperature (EXT, EREF, DD5, FFP, MAT, MCMT, and MWMT). For example, EREF (Hargreave’s reference evaporation), which represents potential evaporation, is calculated using both temperature and radiation data (Aguilar and Polo 2011). These results emphasize that conclusions about which climate variables explain variation in genecological models depend on which environments are sampled. In the PNW, moisture variables are expected to be better predictors of genecological variation and field performance in region-wide experiments, whereas temperature variables will be better predictors locally (e.g., St.Clair et al. 2019), particularly if diverse elevations are sampled. Thus, for region-wide evaluation of seed deployment, we used a multivariate climate distance function that includes multiple climate variables associated with temperature and moisture.

Our final set of climate variables accounts for the diverse effects of climate on forest trees. MAT and FFP account for differences in annual temperatures, growing season lengths, and frost events, which have been associated with population differences in tree productivity (Wang et al. 2006, Wang et al. 2010, Gray et al. 2011, Cortini et al. 2012) and cold adaptation (Howe et al. 2003, Prevéy et al. 2018). MWMT accounts for genecological variation associated with summer maximum temperatures (Parker and van Niejenhuis 1996) and was associated with climate related growth in lodgepole pine and red alder (*Alnus rubra* Bong.) (Hamann and Wang 2005, Cortini et al. 2012). EMT and MCMT account for adaptive variation associated with winter minimum temperatures (Parker and van Niejenhuis 1996, Ukrainetz et al. 2011, St.Clair et al. 2019), and related variables were used to help construct provisional seed zones for the continental U.S. (Bower et al. 2014). Temperature differences between mild coastal sites versus inland sites are reflected in TD, also known as continentality (TD = MWMT ‒ MCMT). TD and related variables have been associated with productivity of European silver fir (Hlásny et al. 2017) and survival and resistance to Rhabdocline needle cast disease in Douglas-fir (Wilhelmi et al. 2017, St.Clair et al. 2019). EREF, MSP, and MAP are important moisture-related variables, and EREF accounts for the effects of evaporative demand (Liu and El-Kassaby 2018). In Douglas-fir, MSP and MAP were associated with population differences in Rhabdocline disease and flowering time (Wilhelmi et al. 2017, Prevéy et al. 2018).

After identifying important climate variables, we developed and studied the performance of a Euclidean climate distance function. Similar measures of climate distance have been developed using some of the same climate variables. For example, O’Neill et al. (2017) used seven ClimateWNA variables plus latitude to guide seed transfer in BC, whereas Hamann et al. (2015) used nine ClimateWNA variables to study the speed at which species or populations would need to migrate to track climate change. Using nine climate variables, we identified matching zones using the normalized Euclidean distance among the marginal medians (i.e., vectors of univariate medians) of all target zones. The standardized Euclidean distance has been used to guide climate-based seed transfer in forest trees and other plants (Rehfeldt et al. 2014, Doherty et al. 2017, O’Neill et al. 2017, Shryock et al. 2018), and was judged to be suitable for identifying climate analogs based on statistical considerations and simulations (Grenier et al. 2013). Rather than using a multivariate climate distance (i.e., single metric), multiple threshold approaches have also been used. Hamann et al. (2015), for example, used independent thresholds for two multivariate climate variables (e.g., principal component scores, PC1 and PC2) to find matching climates. The Seedlot Selection Tool (US-SST) uses a threshold approach as the default method for finding matching climates (St.Clair et al. 2022). First, an acceptable deployment area is found by excluding all locations with distances greater than the CDT for any of the climate variables. Then, a multivariate Euclidean climate distance is calculated for the remaining locations (St.Clair et al. 2022). Although independent thresholds may be appropriate for PC scores, they could be too conservative for individual climate variables. As more independent variables are added, the deployment area shrinks. Moreover, the resulting matches tend to be overly influenced by the climate variable with the most restrictive CDT. Thus, the threshold approach can produce misleading results, particularly if one includes an overly restrictive CDT for an unimportant climate variable.

In some cases, climate principal components, rather than original climate variables, were used to calculate climate distances (Hamann et al. 2015, Shryock et al. 2018). This standardizes the variables, eliminates multicollinearity, and weights the variables according to their associations with major climatic gradients (Shryock et al. 2018). However, it is challenging to decide how to incorporate multiple PC scores into a single index of climate distance. Instead, we used the original climate variables (normalized), after removing highly correlated variables. This provides a foundation for incorporating explicit climate variable weights in the future, which may be able to account for species’ individualized responses to climate.

When we compared the performance of our 9-variable climate distance function to functions ranging from 1 to 16 variables, the univariate distance functions yielded more matches for each climate variable (data not shown). Thus, they were less conservative than the multivariate functions shown in Figure 6a. We also found differences among the univariate distance functions, which can be explained by their different patterns of variation (e.g., small-scale versus large-scale variation). Using TD, for example, we found substantially more matches than we did using MAT, presumably because of their different patterns of geographic variation. Differences among important climate variables emphasize the need for a multivariate distance function to balance their different patterns of variation. However, if redundant or unimportant variables are included, these distance functions may become inappropriate. Based on our random forest analyses, the classification and regression performance of the distance functions plateaued after eight to nine climate variables (data not shown).

### Climate distance thresholds (CDTs)

Because there are few provenance tests spanning our study area, we inferred CDTs from within-zone climate variation. The CDTs we calculated (RANGE/2) are approximately equal to the average transfer distances within a zone. The resulting CDTs averaged 0.66 for the geographic zone sets and 1.10 for the ecological zone sets (Appendix S1: Table S9). To allow easy interpretation of climate distances, Zone Matcher reports absolute climate distances (CD) and scaled climate distances, where the scaled climate distance equals CD/CDT. CDTs for the geographic zone sets were similar, ranging from 0.60 for OR66 to 0.71 for WA66 (Appendix S1: Table S9). The CDT for the EPA4 zone set (1.16) was larger than for BEC (1.04), suggesting the BEC zones are more climatically homogeneous than are the EPA4 zones (Appendix S1: Table S9).

In contrast to our approach, the province of BC infers ‘critical seed transfer distances’ (CSTDs) from regional provenance tests in BC using an approach similar to that described by O’Neill et al. (2017). Using updated climate transfer functions for eight conifer species (Greg O’Neill, unpublished), we converted the BC CSTDs to CDTs that are based on our 9-variable climate distance function. The resulting CDTs ranged from 0.807 for lodgepole pine to 1.380 for Engelmann spruce. Based on the BC values and the CDT values we reported above, we included three CDT options in Zone Matcher: Conservative = 0.7, Moderate = 0.9, and Liberal = 1.1.

Because we used zone marginal medians to calculate inter-zone distances, only about 25% to 30% of the distances between individual sites were predicted to be less than the CDT (Figure 7). Instead of being hard cutoffs, the CDTs were designed to be the smallest distances that might result in detectable changes in tree growth. For example, about 14% of the within-zone distances for the WA66 zone set (i.e., inter-zone distance = 0) exceeded the CDT of 0.7. This is relevant because the calculated CDT for this zone set was 0.71 and the WA66 zones are considered overly conservative. Although shorter climate distances should be given preference, transfers between zones with a CDT of 1.1 are probably reasonable, particularly when other options are unavailable. For example, the CDTs estimated from provenance tests, which ranged from 0.807 to 1.380, were based on predicted height reductions of only 2.5%. Overall, the CDTs are thresholds beyond which climate transfers *may* have observable effects. New provenance tests will be important for refining CDTs, particularly for areas outside of BC and for new species.

### Characteristics of geographic and ecological zones

To select the base zones for Zone Matcher, we considered which zone sets are widely used, the type of zone set (i.e., geographic versus ecological), and zone size. Rather than designing a new seed deployment system, we based our focal zone system on existing, widely used zone sets.

OR66, WA66, OR96, and WA02 are generic and species-specific zone sets widely used in the PNW (DeBell 2019, Oregon Department of Forestry 2021). The CA zone set is the only geographic zone set that spans the entire state of California (Buck et al. 1970). The original ID/MT zone set, used by the Inland Empire Tree Improvement Cooperative, consisted of a gridded system of zone units with 100-foot elevation bands. We adjusted these zones to have elevation bands of 500-feet to be more consistent with bands for the rest of the region.

The BEC zones are variant-level zones from the British Columbia Biogeoclimatic Ecosystem Classification system (DeLong et al. 2010, MacKenzie and Meidinger 2018). These ecological zones have been used for seed deployment in BC since 2018, and the former geographic zone system has been phased out (Nicholls 2019). EPA Level IV (EPA4) ecoregions are widely used for research and forest management in the U.S., including for non-forest seed transfer (Omernik and Gallant 1986, Miller et al. 2011, Bower et al. 2014). On average, the BEC and EPA4 zones are much larger (365 K ha) than the geographic zones (24 K ha), which is an advantage when they are used as fixed zones. When they are used as focal zones, ecological zones can be evaluated based on similarity of climate plus other ecological factors, whereas matches among geographic zones are based on climate alone. Although EPA4 zones are included in Zone Matcher, they have not been widely used to guide seed deployment. Thus, unlike the other zone sets, they have not been tested empirically. Because of their larger size and greater climate variability, CDTs inferred from ecological zones are about 1.7 times as large as CDTs inferred from geographic zones.

These geographic and ecological zone sets cover the entire PNW region and are used by most major public and private forest landowners. Thus, forest managers will be able to adopt our new focal zone system with little adverse impact on existing procedures for seed collection, breeding, seed and seedling production, and reforestation planning.

### How well do existing zones control maladaptation?

Many, small deployment zones are better for controlling maladaptation but are more costly (O’Neill and Aitken 2004, Ukrainetz et al. 2011, Doherty et al. 2017). Our analyses of climate variation allowed us to judge whether existing zones are small enough to prevent maladaptation. Seed zones and breeding zones have gradually expanded over time (Randall 1996, Stonecypher 1996, Johnson 1997, Randall and Berrang 2002). Thus, older geographic seed zones, such as OR66 and WA66, are likely smaller than necessary to control maladaptation. Furthermore, the geographic zones are much smaller than the ecological zones.

We used CDT_ABS_ for the individual climate variables to assess whether existing zones control maladaptation. CDT_ABS_ for the geographic zone sets were generally lower than acceptable transfer distances inferred from genecological studies. For MAT, CDT_ABS_ values were mostly below the 2-3°C values recommended from provenance test results (Wang et al. 2006, St.Clair et al. 2019) (Appendix S1: Figure S9). For the generic (non-species specific) zone sets, CDT_ABS_ for MAT ranged from 0.75°C for OR66 to 2.20°C for EPA4, and for the Douglas-fir zone sets, they were 1.15°C (Figure 3, Appendix 1: Table S9). For MSP, Wilhelmi et al. (2017) concluded that a CDT_ABS_ of 159 mm MSP was necessary to avoid unacceptable levels of Rhabdocline infection, especially for transfers from low to high MSP. However, we know of no studies that have estimated MSP CDT_ABS_ for other traits. CDT_ABS_ for MSP ranged from 20.5 mm for the CA zone set to 79.7 mm for BEC and 106.1 mm for the Douglas-fir zone sets. Although precipitation CDT_ABS_ should be smaller for drier regions, these CDTs seem to be well below what is needed to avoid excessive maladaptation, at least for MAT and MSP. In contrast, maximum within-zone transfer distances (i.e., RANGE = 2 × CDT_ABS_) exceeded recommended values for MAT for the EPA4 zone set (4.4°C). Excluding EPA4, the largest MAT RANGE value was 2.94°C for BEC, and the smallest value was 1.50°C for OR66.

Using Zone Matcher’s focal zone system, the base zones should be small enough for the most risk-averse users to feel confident about controlling maladaptation. Then, CDTs can be adjusted for higher risk tolerance or to accommodate differences among individual tree species. Although differences in the extent of local adaptation are often assumed to exist among species (Rehfeldt 1994, Randall and Berrang 2002, Johnson et al. 2004), they have been rarely documented in direct comparative studies or using identical analytical approaches (but see Frank et al. 2017). In any case, to accommodate differences in local adaptation among species, it is simpler to modify the CDTs, rather than the zones themselves. This approach allows a single set of generic zones to be used for all species.

### Climate-based seed deployment and assisted migration

Land managers must consider various sources of uncertainty in making reforestation decisions. For example, climate projections are affected by uncertainty in future population sizes, policies, and technologies to mitigate climate change. Zone Matcher relies on climate data from ClimateNA. Future projections from ClimateNA v7.42 are based on CMIP6 data and an ensemble of eight climate models (Mahony et al. 2022). In addition to the uncertainty of future climates, climate interpolation models may not accurately depict climate variation among sites, particularly in the PNW region where the density of weather stations is low. Finally, responses to climate change will vary among species.

Seed deployment and assisted migration should not be based on climate alone. Deployment decisions should consider other expert knowledge of the site, such as soil characteristics, microclimatic factors such as aspect and microtopography, and potential for pests and fire. Current and future modeled species distributions can be used to avoid deploying seed to areas where the species’ habitat is unlikely to occur. Information on projected species distributions for eight key species is available in Zone Matcher.

Although provenance tests can be used to infer changes in growth or productivity from climate change (e.g., Wang et al. 2006), Zone Matcher does not report these values because of large uncertainties. First, there are no genetic tests that adequately sample the full Zone Matcher region. Because climate regimes vary regionally and at different scales, genecological models developed in one region may be inappropriate in others. Second, even without extrapolation, predictions of growth and survival from genecological models have large errors (see figures in Wang et al. 2010, Ukrainetz et al. 2011, St.Clair et al. 2019), particularly using small numbers of provenances or plantations (Zhao et al. 2023). Third, relative performance changes over time, including changes in rank among provenances (Clausen 1973, Zas et al. 2004, St.Clair et al. 2019), perhaps because rare climatic events affect some provenances more than others. Fourth, climate interpolation models contain error (Ye et al. 2022) and projections of future climates are highly uncertain (Lehner et al. 2023, Yoshikawa et al. 2023, Evin et al. 2024). Finally, genecological models mostly assume the climates where trees are deployed will be static, yet new plantations will probably experience substantial changes in climate over time. Overall, the propagation of these errors makes future projections of forest ecosystem responses highly uncertain as well (Yoshikawa et al. 2023).

### Other seed deployment tools

Two other web-based tools for deploying tree seed have been described in the literature: the CBST Seedlot Selection Tool used in BC (O’Neill et al. 2017) and the U.S. Seedlot Selection Tool that covers all of North America (St.Clair et al. 2022). The CBST Seedlot Selection Tool (BC-SST; https://maps.forsite.ca/204/SeedTransfer/) uses a focal zone approach to guide seed deployment in BC. However, it differs from Zone Matcher in four main ways. First, because the BC-SST zones (BEC zones) do not extend beyond BC, it cannot be used to easily deploy BC seedlots outside of BC or find seedlots from elsewhere to deploy in BC. Second, the BC-SST uses a Euclidean distance function that includes a different set of ClimateNA variables plus latitude, which is used to account for differences in photoperiod (O’Neill et al. 2017). The Zone Matcher distance function is based on climate variables alone, and rather than latitude, Zone Matcher reports photoperiod directly for potential seed transfers. Nonetheless, the Zone Matcher and BC-SST distance functions were highly correlated (r = 0.99). Third, because species differ in their response to a given climate transfer distance, the BC-SST uses species-specific ‘genetic suitabilities’ to assess potential seed transfers. These suitabilities are estimates of relative height growth for deployed seedlots compared to local seedlots based on large provenance tests in BC (e.g., O’Neill et al. 2017). Genetic suitabilities have been calculated for ten species: Douglas-fir, western larch, lodgepole pine, ponderosa pine, interior spruce (*Picea engelmanii* × *glauca*), grand fir, western redcedar, western hemlock, western white pine, and Alaska-cedar (*Callitropsis nootkatensis* (D. Don) Oerst.). Fourth, the BC-SST uses a single CDT and assisted migration scenario to evaluate deployment options. In Zone Matcher, the use of assisted migration is optional but can be implemented using 5 historic climates and 12 future climate scenarios.

The U.S. Seedlot Selection Tool (US-SST) can be used to guide seed deployment across North America (St.Clair et al. 2022). Unlike Zone Matcher and the BC-SST, the US-SST uses a focal point system. Thus, there is no direct link between the mapped output and seed deployment zones. Nonetheless, the US-SST base map can be queried to see which zones occur at a particular location. Despite that capability, the US-SST does not have zones for BC, Idaho/Montana, EPA4 ecoregions, or other zone sets found in Zone Matcher. A potential advantage of the US-SST is the high resolution of the maps (∼0.5 km), but this may be misleading because of uncertain climate predictions (discussed above) and large genetic (seedlot) sampling errors. Genetic sampling errors are large because 50% or more of the total genetic variation for climate adaptation traits typically occurs within populations (Howe et al. 2003, Savolainen et al. 2007, Cooper et al. 2022). Finally, most planted seedlots do not have precise climatic origins or known deployment areas. These include bulked seedlots from native stands and orchard seedlots, particularly when pollen contamination is present. The US-SST also differs in other ways. Zone Matcher and the BC-SST use a single multivariate climate distance function, whereas US-SST users can choose their own set of user-defined climate variables which are used in a threshold matching approach (discussed above). However, users can choose to use genecological functions for lodgepole pine or Engelmann spruce instead. Finally, Zone Matcher uses climate distance functions derived from normalized climate variables and data from the CMIP6 project for future climate projections (Mahony et al. 2022). Currently, the US-SST uses non-normalized climate variables from the CMIP5 project (St.Clair et al. 2022), but plans are underway to update the climate data to CMIP6 (Vicky Erickson, personal communication).

### Recommendations and next steps

Zone Matcher can be used to find seedlots for a planting site, or planting sites for a seedlot. To find seedlots, the focal zone is the zone where the seedlings will be planted. Then, Zone Matcher returns a ranked list of target zones that correspond to acceptable seedlots, which are seedlots collected from the target zones or seedlots that can be deployed there (e.g., for orchard seed). To find planting sites for a seedlot, the focal zone is the zone where the wild-stand or orchard seedlot is expected to perform best. Then, Zone Matcher returns a ranked list of planting zones. In either case, users should pick the closest match based on constraints such as the availability of seeds or seedlings. An analogous approach can be used for deploying trees produced by vegetative propagation.

To find climate matches, the user must choose climate scenarios for the seedlot and planting site. Because of recent climate change, populations may already be maladapted to current climates (Gray et al. 2011, Gray and Hamann 2013, Aitken and Bemmels 2016). Thus, native seedlots are probably best adapted to recent historical climates, which, depending on species and stand history, might include climate normals from 1941-1970 or 1961-1990 (O’Neill et al. 2017, O’Neill and Gómez-Pineda 2021, St.Clair et al. 2022). Assisted migration will mostly involve moving populations from warmer to colder environments. To accomplish this, a target year and future climate scenario must be chosen for the planting site. The target year should be distant enough that trees remain well-adapted over the long-term (e.g., many decades), while avoiding cold damage at the vulnerable seedling stage (St.Clair and Howe 2007). For managed forests, recommended target years are when the plantation is one-quarter to one-third rotation age (Ukrainetz et al. 2011, O’Neill et al. 2017, O’Neill and Gómez-Pineda 2021). Next, the SSP can be chosen based on the user’s risk tolerance or assumptions about the pace of climate change. A consensus approach is to select zones that match multiple climate scenarios (i.e., target years and SSPs).

Overall, we recommend users favor closer climatic matches over more distant ones, and use large CDTs with caution (e.g., CDT > 1.1). Further research is needed to refine estimates of climate distance thresholds across species and subregions. New provenance tests should be established, and existing provenance tests should be used to quantify the relationship between field performance versus Zone Matcher climate distance. Because of the many uncertainties discussed above, we designed Zone Matcher to match seedlots and planting sites based on climate, rather than predict plantation performance. Finally, before selecting a seedlot or planting site, users should check the differences in the individual climate variables between the focal and target zones. At least for MAT, these values can be compared to the CDT_ABS_ of 2 to 3°C that have been inferred from provenance test results (Wang et al. 2006, St.Clair et al. 2019) (discussed above).

The use of climate matching to guide seed deployment rests almost entirely on research with natural populations. Yet, the majority of trees planted in the Zone Matcher region come from seed orchards. Many of these orchards consist of parents from diverse locations or their advance generation progeny. Additionally, because of pollen contamination, it is common for 20% to 40% of orchard seeds to have been fathered by local trees outside the orchard (Slavov et al. 2005). Thus, orchard seedlots do not have precise geographic origins in the same sense as do seedlots collected from the wild. Instead, orchard seedlots are usually associated with breeding zones that have been delineated and validated using field progeny tests. In general, these deployment zones can be analyzed easily using Zone Matcher. We are currently developing a new tool to improve delineation of orchard deployment zones by considering the geographic origins of the first-generation parents, pedigree of the orchard clones, number of ramets per clone, and extent of local pollen contamination.

## DATA AVAILABILITY STATEMENT

Raw data and metadata used to generate tables and figures, novel code utilized to generate results, and derived data products have been archived on Dryad: https://doi.org/10.5061/dryad.931zcrjww.

## ACKNOWLEDGEMENTS

We thank Scott Kolpak (USFS) and Vicky J. Erikson (USFS) for providing zone shapefiles, and Marc Rust (Inland Empire Tree Improvement Cooperative) for helping to produce the IETIC zones. We thank members of the Pacific Northwest Tree Improvement Research Cooperative for guidance and financial support and The Northwest Climate Adaptation Science Center (NW CASC-11700) for additional funding.

## AUTHOR CONTRIBUTIONS

TJS conducted analyses, wrote text, and produced figures; MLM conducted analyses, wrote text, and produced figures; NS-M helped construct the DEM, zone shapefiles, zone rasters, and initial climate data; GAO provided the provenance transfer models and critical seed transfer distances used to infer climate distance thresholds in BC; TW provided the species distribution models; JAES wrote code for the Shiny web application; and GTH conceived the project, conducted analyses, wrote text, produced figures, and designed the shiny web application. All authors reviewed and edited the manuscript.

