## Appendix S1 for "Zone Matcher: A climate-based web application for deployment and assisted migration of forest trees"

**Journal: Ecological Applications**

### Tables

**Table S1. Tree deployment zones in the Pacific Northwest.** Species are listed as their USDA plant symbol (USDA PLANTS Database).

| Abbreviation | Zone coverage | Organization* | Year | Species† | Data source | References |
|---|---|---|---|---|---|---|
| **(a) Ecological zone sets** | | | | | | |
| BEC | British Columbia | BCMoF | 2021 | All (generic) | Ministry of Forests - Forest Analysis and Inventory. (2021). *BEC Map (version 12)*. Retrieved from https://catalogue.data.gov.bc.ca/dataset/bec-map | Meidinger and Pojar (1991) |
| EPA4 | EPA Level IV Ecoregions | EPA | 1986 | All (generic) | United States Environmental Protection Agency. *US Level IV Ecoregions.* Retrieved from https://www.epa.gov/eco-research/level-iii-and-iv-ecoregions-continental-united-states | Omernik and Gallant (1986) |
| **(b) Geographic zone sets** | | | | | | |
| CA | California | USFS | 1970 | All (generic) | Fire and Resource Assessment Program (Calif.). *California Tree Seed Zones Map.* Retrieved from https://maps.princeton.edu/catalog/stanford-sg575tf7838 | Buck et al. (1970) |
| ID/MT | Idaho, E. Washington, W. Montana | IETIC | 2020 | All (generic) | Developed for this paper | |

| | | | | | | |
|---|---|---|---|---|---|---|
| OR96 | Oregon | ODF | 1996 | ABAM, ABCO, ABGR, ABMAS, ABPR, ALRU2, CADE27, CANO9, CHLA, PICO, PICOC, PIEN, PIJE, PILA, PIMO3, PIPO, PISI, POBAT, PSME, TABR2, THPL, TSHE, and 10 generic zones‡ | Oregon Department of Forestry. *Seed Zones*. Retrieved from https://www.oregon.gov/ODF/AboutODF/Pages/MapsData.aspx | Sorensen (1979); Randall (1996) |
| OR66 | Oregon | WFTSC | 1966 | All (generic) | Vicky Erickson, USFS Region 6 Geneticist | Randall (1996) |
| WA66 | Washington | WFTSC | 1966 | All (generic) | Vicky Erickson, USFS Region 6 Geneticist | Randall (1996) |
| WA02 | Washington | WDNR | 2002 | ABAM, ABGR, ABPR, ALRU2, CANO9, LAOC, PICO, PIEN, PIMO3, PIPO, PISI, POBAT, PSME, TABR2, THPL, and TSHE‡ | Vicky Erickson, USFS Region 6 Geneticist | Randall and Berrang (2002) |

*BCMoF is British Columbia Ministry of Forests, EPA is U.S. Environmental Protection Agency, IETIC is Inland Empire Tree Improvement Cooperative, ODF is Oregon Department of Forestry, USFS is United States Forest Service, WFTSC is Western Forest Tree Seed Council, and WDNR is Washington Department of Natural Resources.

[†]ABAM is Pacific silver fir, ABCO is white fir, ABGR is grand fir, ABMAS is Shasta fir, ABPR is noble fir, ALRU2 is red alder, CADE27 is incense-cedar, CANO9 is Alaska yellow cedar, CHLA is Port-Orford-cedar, LAOC is western larch, PICO is lodgepole pine, PICOC is shore pine, PIEN is Engelmann spruce, PIJE is Jeffrey pine, PILA is sugar pine, PIMO3 is western white pine, PIPO is ponderosa pine, PISI is Sitka spruce, POBAT is black cottonwood, PSME is Douglas-fir, TABR2 is Pacific yew, THPL is western redcedar, TSHE is western hemlock, and TSME is mountain hemlock.

[‡]For OR96 and WA02, only species-specific zones for which shapefiles existed were analyzed. This resulted in five species-specific zones being analyzed for OR96 and WA02.

**Table S2. Climate variable abbreviations, definitions, and units.**

| Abbreviation | ClimateNA variable | Units |
|---|---|---|
| AHM | Annual heat-moisture index (MAT+10)/(MAP/1000) | °C/m |
| CMD | Hargreaves climatic moisture deficit | mm |
| DD_0 | Degree-days below 0˚C, chilling degree-days | dd |
| DD5 | Degree days above 5˚C, growing degree-days | dd |
| EMT | Extreme minimum temperature over 30 years | °C |
| Eref | Hargreaves reference evaporation | mm |
| EXT | Extreme maximum temperature over 30 years | °C |
| FFP | Frost-free period | days |
| MAP | Mean annual precipitation | mm |
| MAT | Mean annual temperature | °C |
| MCMT | Mean coldest month temperature | °C |
| MSP | Mean summer precipitation [May to September] | mm |
| MWMT | Mean warmest month temperature | °C |
| PAS | Precipitation as snow [August to July] | mm |
| SHM | Summer heat-moisture index (MWMT)/(MSP/1000) | °C/m |
| TD | Temperature difference between MWMT and MCMT, or continentality | °C |

**Table S3. Zone set characteristics for generic (non-species-specific) forested zones in the Pacific Northwest.** Values are sums (numbers of zone units and zones) or means (sizes and elevations) for a given zone set.

| Zone set | | Zone unit | | | | Zone | | | | | |
|---|---|---|---|---|---|---|---|---|---|---|---|
| ID* | Species† | No. of zone units | Size (ha)‡ | Elevation lower limit (ft)‡ | Elevation upper limit (ft)‡ | No. of zones | No. of zones/ unit | Size (ha)‡ | Elevation lower limit (ft)‡ | Elevation upper limit (ft)‡ | Elevation band width (ft)‡ |
| **(a) Ecological zone sets** | | | | | | | | | | | |
| BEC | GENERIC | 200 | 407,663 | 2,414 | 5,383 | 200 | 1 | 407,663 | 2,414 | 5,383 | 2,970 |
| EPA4 | GENERIC | 253 | 322,656 | 2,118 | 6,168 | 253 | 1 | 322,656 | 2,118 | 6,168 | 4,049 |
| Mean | | 227 | 365,160 | 2,266 | 5,776 | 227 | 1 | 365,160 | 2,266 | 5,776 | 3,509 |
| **(b) Geographic zone sets** | | | | | | | | | | | |
| CA | GENERIC | 84 | 466,184 | 1,147 | 7,541 | 900 | 11 | 28,780 | 4,526 | 4,991 | 465 |
| ID/MT | GENERIC | 209 | 215,375 | 3,503 | 8,566 | 1,807 | 9 | 19,115 | 5,754 | 6,216 | 461 |
| OR66§ | GENERIC | 80 | 351,706 | 1,797 | 6,638 | 731 | 9 | 22,726 | 4,084 | 4,536 | 452 |
| WA66 | GENERIC | 52 | 343,177 | 673 | 7,153 | 555 | 11 | 24,017 | 3,264 | 3,729 | 465 |
| Mean | | 106 | 344,110 | 1,780 | 7,474 | 998 | 10 | 23,660 | 4,407 | 4,868 | 461 |

*BEC is British Columbia's Biogeoclimatic Ecosystem Classification; CA is California; EPA4 is U.S. Environmental Protection Agency Level IV Ecoregions; ID/MT is Inland Empire Tree Improvement Cooperative zones in Idaho/western Montana; OR66 is Oregon's seed zones published in 1966; WA66 is Washington's seed zones published in 1966. OR96 is Oregon's seed zones published in 1996 and WA02 is Washington's seed zones published in 2002 (see Table S1).

†GENERIC indicates the zone set is not species specific.

‡Because sizes and elevations are based on raster sampling, they differ slightly from the original design of the zones but correspond exactly to sampled climate data.

§The OR66 data includes zone units and zones duplicated in the WA66 zone set. Duplicated zones were omitted from statistical analyses.

**Table S4. Zone set characteristics for species-specific forested zones in the Pacific Northwest.** Values are sums (numbers of zone units and zones) or means (sizes and elevations) for a given zone set.

| Zone set | | Zone unit | | | | Zone | | | | | |
|---|---|---|---|---|---|---|---|---|---|---|---|
| ID* | Species[†] | No. of zone units | Size (ha)[‡] | Elevation lower limit (ft)[‡] | Elevation upper limit (ft)[‡] | No. of zones | No. of zones/unit | Size (ha)[‡] | Elevation lower limit (ft)[‡] | Elevation upper limit (ft)[‡] | Elevation band width (ft)[‡] |
| **(a) Geographic species-specific zone sets** | | | | | | | | | | | |
| OR96 | PICO | 7 | 859,372 | 2,435 | 8,696 | 39 | 6 | 150,264 | 4,571 | 5,487 | 916 |
| OR96 | PIMO | 5 | 1,282,809 | 173 | 8,272 | 5 | 1 | 1,282,809 | 173 | 8,272 | 8,099 |
| OR96 | PIPO | 15 | 644,089 | 1,938 | 7,827 | 86 | 6 | 107,002 | 4,361 | 5,232 | 871 |
| OR96 | PSME | 16 | 484,424 | 552 | 6,502 | 163 | 10 | 47,517 | 3,391 | 3,903 | 512 |
| OR96 | THPL | 4 | 1,937,945 | 468 | 7,478 | 6 | 2 | 1,291,958 | 1,481 | 6,151 | 4,670 |
| WA02 | PICO | 17 | 457,069 | 649 | 7,291 | 109 | 6 | 70,931 | 3,048 | 3,862 | 814 |
| WA02 | PIMO | 7 | 1,143,784 | 653 | 8,624 | 7 | 1 | 1,143,784 | 653 | 8,624 | 7,971 |
| WA02 | PIPO | 11 | 506,582 | 1,169 | 7,246 | 58 | 5 | 95,962 | 3,434 | 4,476 | 1,042 |
| WA02 | PSME | 16 | 676,337 | 502 | 7,758 | 106 | 7 | 101,311 | 2,982 | 3,815 | 833 |
| WA02 | THPL | 7 | 1,200,402 | 584 | 8,802 | 29 | 4 | 289,010 | 2,884 | 4,456 | 1,572 |
| Mean | | 11 | 919,281 | 912 | 7,850 | 61 | 5 | 458,055 | 2,698 | 5,428 | 2,730 |

*See Table S3 for a description of the zone set IDs.

[†]See Table S1 for a description of species names from USDA plant symbols (USDA PLANTS database).

[‡]Because sizes and elevations are based on raster sampling, they differ slightly from the original design of the zones but correspond exactly to sampled climate data.

**Table S5. Correlations of geographic and climate variables for four non-overlapping geographic zone sets in the Pacific Northwest (CA, ID/MT, OR66, and WA66).** The upper diagonal shows the average of the correlations for each zone set. The lower diagonal shows the correlations calculated across all four zone sets. Correlations ≥ |0.95| are in bold.

| | LAT | LONG | PHOTO | AHM | CMD | DD_0 | DD5 | EMT | EREF | EXT | FFP | MAP | MAT | MCMT | MSP | MWMT | PAS | SHM | TD |
|---|---|---|---|---|---|---|---|---|---|---|---|---|---|---|---|---|---|---|---|
| LAT | – | -0.09 | **1.00** | -0.15† | -0.30 | 0.11 | -0.07† | -0.06 | -0.19 | -0.05† | 0.00† | 0.16† | -0.09 | -0.15 | 0.37† | -0.05† | 0.13† | -0.33† | 0.12 |
| LONG | 0.27 | – | -0.10 | 0.42† | 0.23 | 0.46 | -0.18† | -0.57 | -0.08 | -0.01† | -0.37 | -0.57 | -0.29† | -0.50 | -0.30 | -0.02† | -0.01† | 0.21 | 0.54 |
| PHOTO | **1.00** | 0.27 | – | -0.16† | -0.30 | 0.11 | -0.07† | -0.06 | -0.19 | -0.05† | 0.00† | 0.16† | -0.09 | -0.15 | 0.37† | -0.06† | 0.13† | -0.32† | 0.12 |
| AHM | -0.29 | 0.21 | -0.30 | – | 0.91 | -0.31† | 0.59 | 0.10† | 0.67 | 0.71 | 0.32† | -0.95 | 0.49 | 0.22† | -0.86 | 0.69 | -0.75 | 0.90 | 0.56 |
| CMD | -0.64 | -0.22 | -0.64 | 0.815 | – | -0.47 | 0.71 | 0.26† | 0.82 | 0.84 | 0.41 | -0.80 | 0.63 | 0.39† | -0.88 | 0.80 | -0.78 | 0.95 | 0.48† |
| DD_0 | 0.49 | 0.70 | 0.50 | -0.33 | -0.67 | – | -0.90 | **-0.95** | -0.82 | -0.74 | -0.93 | 0.07 | **-0.96** | **-0.99** | 0.22† | -0.77 | 0.78 | -0.44 | 0.27 |
| DD5 | -0.43 | -0.43 | -0.44 | 0.59 | 0.81 | -0.91 | – | 0.78 | 0.94 | 0.93 | 0.90 | -0.37 | **0.98** | 0.83 | -0.45 | **0.96** | -0.90 | 0.66 | 0.11 |
| EMT | -0.42 | -0.77 | -0.44 | 0.17 | 0.52 | **-0.97** | 0.83 | – | 0.63 | 0.56 | 0.91 | 0.14 | 0.86 | **0.97** | -0.06 | 0.62 | -0.62 | 0.26† | -0.45 |
| EREF | -0.56 | -0.37 | -0.57 | 0.65 | 0.90 | -0.86 | **0.95** | 0.73 | – | **0.96** | 0.73 | -0.47 | 0.91 | 0.74 | -0.54 | 0.94 | -0.88 | 0.73 | 0.21† |
| EXT | -0.33 | -0.24 | -0.34 | 0.70 | 0.85 | -0.75 | 0.93 | 0.62 | 0.94 | – | 0.70 | -0.52 | 0.87 | 0.65 | -0.56 | **0.96** | -0.87 | 0.74 | 0.34† |
| FFP | -0.34 | -0.62 | -0.36 | 0.33 | 0.59 | -0.94 | 0.91 | 0.94 | 0.77 | 0.72 | – | -0.09† | 0.93 | 0.90 | -0.22 | 0.78 | -0.76 | 0.43 | -0.18 |
| MAP | 0.14 | -0.48 | 0.15 | -0.92 | -0.60 | -0.02† | -0.27 | 0.18 | -0.36 | -0.44 | 0.00† | – | -0.26† | 0.01 | 0.83 | -0.50 | 0.60 | -0.78 | -0.64 |
| MAT | -0.46 | -0.56 | -0.48 | 0.49 | 0.77 | **-0.97** | **0.98** | 0.90 | 0.93 | 0.87 | 0.94 | -0.15 | – | 0.91 | -0.37† | 0.90 | -0.86 | 0.59 | -0.04† |
| MCMT | -0.53 | -0.74 | -0.54 | 0.27 | 0.64 | **-0.99** | 0.87 | **0.98** | 0.82 | 0.69 | 0.92 | 0.07 | 0.94 | – | -0.16 | 0.69 | -0.70 | 0.37† | -0.39† |
| MSP | 0.72 | 0.25 | 0.73 | -0.74 | -0.92 | 0.57 | -0.65 | -0.47 | -0.74 | -0.63 | -0.50 | 0.57 | -0.64 | -0.56 | – | -0.54 | 0.60 | -0.95 | -0.43 |
| MWMT | -0.38 | -0.26 | -0.39 | 0.69 | 0.85 | -0.79 | **0.96** | 0.68 | 0.94 | **0.96** | 0.80 | -0.42 | 0.91 | 0.73 | -0.66 | – | -0.87 | 0.73 | 0.35 |
| PAS | 0.45 | 0.23 | 0.46 | -0.75 | -0.84 | 0.79 | -0.91 | -0.67 | -0.90 | -0.87 | -0.76 | 0.51 | -0.86 | -0.73 | 0.71 | -0.88 | – | -0.74 | -0.15† |
| SHM | -0.68 | -0.31 | -0.69 | 0.76 | **0.96** | -0.70 | 0.79 | 0.58 | 0.85 | 0.76 | 0.64 | -0.53 | 0.77 | 0.68 | **-0.97** | 0.79 | -0.80 | – | 0.41 |
| TD | 0.41 | 0.77 | 0.43 | 0.34 | -0.03 | 0.60 | -0.25 | -0.71 | -0.20 | 0.02† | -0.49 | -0.56 | -0.41 | -0.68 | 0.13 | -0.01† | 0.15 | -0.15 | – |

*Climate variables are described in Table S2.

†Values are non-significant at the 0.05 level (2-tailed), all others are statistically significant. *P*-values were adjusted for multiple comparisons using the Holm correction method in the corr.test R function, and *p*-values for the averages (upper diagonal) are averaged across the four zone sets.

**Table S6. Correlations of geographic and climate variables for two non-overlapping ecological zone sets in the Pacific Northwest (BEC and EPA4).** The upper diagonal shows the average of the correlations for each zone set. The lower diagonal shows the correlations calculated across the two zone sets. Correlations ≥ |0.95| are in bold.

| | LAT | LONG | PHOTO | AHM | CMD | DD_0 | DD5 | EMT | EREF | EXT | FFP | MAP | MAT | MCMT | MSP | MWMT | PAS | SHM | TD |
|---|---|---|---|---|---|---|---|---|---|---|---|---|---|---|---|---|---|---|---|
| LAT | -- | -0.20 | **1.00** | -0.37 | -0.58 | 0.58 | -0.52 | -0.53 | -0.65 | -0.52 | -0.47 | 0.26† | -0.60 | -0.61 | 0.61 | -0.46 | 0.49 | -0.62 | 0.41 |
| LONG | -0.33 | -- | -0.20 | 0.23† | -0.01 | 0.38† | -0.17† | -0.45† | -0.04 | 0.04 | -0.35† | -0.38 | -0.24† | -0.40† | 0.00 | -0.02 | 0.02 | -0.07 | 0.51 |
| PHOTO | **1.00** | -0.35 | -- | -0.37 | -0.58 | 0.58 | -0.52 | -0.53 | -0.65 | -0.52 | -0.47 | 0.26† | -0.60 | -0.61 | 0.61 | -0.46 | 0.49 | -0.61 | 0.42 |
| AHM | -0.64 | 0.38 | -0.64 | -- | 0.87 | -0.25† | 0.57 | 0.10† | 0.68 | 0.71 | 0.27† | -0.94 | 0.46 | 0.16† | -0.80 | 0.67 | -0.78 | 0.82 | 0.34† |
| CMD | -0.86 | 0.25 | -0.85 | 0.88 | -- | -0.52 | 0.74 | 0.37† | 0.87 | 0.84 | 0.49 | -0.72 | 0.68 | 0.44† | -0.88 | 0.78 | -0.81 | 0.96 | 0.06 |
| DD_0 | 0.79 | 0.11 | 0.79 | -0.53 | -0.77 | -- | -0.86 | **-0.97** | -0.78 | -0.69 | -0.94 | -0.01† | -0.94 | **-0.98** | 0.37† | -0.72 | 0.64 | -0.54 | 0.62 |
| DD5 | -0.79 | 0.10 | -0.79 | 0.72 | 0.88 | -0.93 | -- | 0.76 | 0.94 | 0.91 | 0.90 | -0.33 | 0.95 | 0.78 | -0.54 | 0.94 | -0.82 | 0.73 | -0.20 |
| EMT | -0.73 | -0.20 | -0.73 | 0.41 | 0.67 | **-0.98** | 0.87 | -- | 0.63 | 0.55 | 0.92 | 0.16† | 0.86 | **0.98** | -0.27† | 0.59 | -0.50† | 0.43† | -0.72 |
| EREF | -0.88 | 0.22 | -0.88 | 0.77 | **0.95** | -0.89 | 0.96 | 0.80 | -- | **0.96** | 0.75 | -0.47 | 0.90 | 0.70 | -0.68 | 0.93 | -0.85 | 0.83 | -0.12 |
| EXT | -0.79 | 0.27 | -0.79 | 0.80 | 0.93 | -0.82 | **0.95** | 0.73 | **0.97** | -- | 0.71 | -0.52 | 0.84 | 0.60 | -0.60 | **0.97** | -0.81 | 0.76 | 0.05 |
| FFP | -0.66 | -0.17 | -0.65 | 0.48 | 0.68 | **-0.95** | 0.91 | **0.95** | 0.81 | 0.77 | -- | -0.01† | 0.93 | 0.90 | -0.33† | 0.77 | -0.62 | 0.53 | -0.47 |
| MAP | 0.40 | -0.44 | 0.40 | -0.91 | -0.66 | 0.22 | -0.44 | -0.10 | -0.51 | -0.56 | -0.18 | -- | -0.21 | 0.09† | 0.73 | -0.47 | 0.65 | -0.66 | -0.49 |
| MAT | -0.80 | 0.02 | -0.80 | 0.65 | 0.85 | **-0.97** | **0.98** | 0.92 | 0.94 | 0.91 | 0.94 | -0.36 | -- | 0.89 | -0.50† | 0.86 | -0.73 | 0.68 | -0.41† |
| MCMT | -0.82 | -0.08 | -0.81 | 0.49 | 0.75 | **-0.99** | 0.90 | **0.98** | 0.87 | 0.79 | 0.93 | -0.17 | **0.95** | -- | -0.33† | 0.62 | -0.54 | 0.48† | -0.73 |
| MSP | 0.82 | -0.16 | 0.82 | -0.85 | -0.92 | 0.71 | -0.78 | -0.64 | -0.85 | -0.78 | -0.63 | 0.69 | -0.76 | -0.69 | -- | -0.54 | 0.78 | -0.94 | 0.02 |
| MWMT | -0.76 | 0.23 | -0.76 | 0.78 | 0.90 | -0.83 | **0.96** | 0.74 | 0.96 | **0.98** | 0.80 | -0.54 | 0.92 | 0.79 | -0.75 | -- | -0.78 | 0.72 | 0.06 |
| PAS | 0.74 | -0.12 | 0.73 | -0.81 | -0.87 | 0.84 | -0.91 | -0.77 | -0.89 | -0.87 | -0.80 | 0.62 | -0.88 | -0.79 | 0.86 | -0.86 | -- | -0.83 | 0.05 |
| SHM | -0.87 | 0.16 | -0.86 | 0.85 | **0.97** | -0.80 | 0.88 | 0.73 | 0.93 | 0.88 | 0.73 | -0.62 | 0.86 | 0.79 | **-0.97** | 0.86 | -0.88 | -- | -0.04 |
| TD | 0.49 | 0.40 | 0.49 | 0.06† | -0.23 | 0.69 | -0.41 | -0.77 | -0.36 | -0.21 | -0.61 | -0.31 | -0.53 | -0.74 | 0.31 | -0.19 | 0.36 | -0.34 | -- |

*Climate variables are described in Table S2.

†Values are non-significant at the 0.05 level (2-tailed), all others are statistically significant. *P*-values were adjusted for multiple comparisons using the Holm correction method in the corr.test R function, and *p*-values for the averages (upper diagonal) are averaged across the two zone sets.

**Table S7. Partitions of climate variation within zone sets.** Variance partitions were averaged among four non-overlapping geographic zone sets (CA, ID/MT, OR66, and WA66) and presented as percent of variance accounted for by zone units, zones within zone units, and within zones.

| Zone unit | | Zone (zone unit) | | Within zone | |
|---|---|---|---|---|---|
| ClimateNA variable* | Variation (%) | ClimateNA variable | Variation (%) | ClimateNA variable* | Variation (%) |
| TD | 67 | EXT | 68 | AHM | 18 |
| EMT | 58 | EREF | 66 | CMD | 17 |
| MSP | 57 | MWMT | 65 | MAP | 17 |
| MAP | 55 | DD5 | 61 | EXT | 17 |
| MCMT | 54 | MAT | 58 | EREF | 17 |
| DD_0 | 47 | PAS | 56 | PAS | 16 |
| SHM | 47 | CMD | 46 | SHM | 16 |
| AHM | 45 | DD_0 | 44 | MWMT | 15 |
| FFP | 45 | FFP | 44 | MSP | 15 |
| CMD | 37 | MCMT | 38 | DD5 | 14 |
| MAT | 30 | SHM | 38 | TD | 13 |
| PAS | 27 | AHM | 36 | MAT | 12 |
| DD5 | 25 | EMT | 35 | FFP | 11 |
| MWMT | 20 | MSP | 28 | DD_0 | 9 |
| EREF | 17 | MAP | 27 | MCMT | 8 |
| EXT | 15 | TD | 20 | EMT | 7 |

*Climate variables are described in Table S2.

**Table S8. Model performance of final climate variables.** Comparison of random forest model performances for four non-overlapping geographic zone sets in the Pacific Northwest (CA, ID/MT, OR66, and WA66) and two non-overlapping ecological zone sets (BEC and EPA4). Classification models were evaluated using OOB error* and regression models were evaluated using RSQ[†].

| | Geographic zone sets | | Ecological zone sets | |
|---|---|---|---|---|
| | Best unique model (8 vars)[‡] | Final common model (9 vars)[§] | Best unique model (8 vars) | Final common model (9 vars) |
| **(a) Classification OOB error*** | | | | |
| | 0.17 | 0.16 | 0.25 | 0.26 |
| **(b) Regression RSQ[†]** | | | | |
| Latitude[‖] | 0.96 | 0.96 | 0.66 | 0.60 |
| Longitude[¶] | 0.87 | 0.90 | 0.36 | 0.34 |

*OOB error is the out-of-bag error associated with the model.

[†]RSQ is the pseudo-R-squared associated with the model (1 - MSE/Var(y)).

[‡]The best unique model is the 8-variate model with the best performance (lowest OOB error or highest RSQ) averaged across six replications.

[§]The final common model is the model performance of the final 9-variate model averaged across six replications.

[‖]Latitude indicates that zone units were sampled across a latitudinal gradient (ranked from southernmost to northernmost) in a sliding window approach.

[¶]Longitude indicates that zone units were sampled across a longitudinal gradient (ranked from westernmost to easternmost) in a sliding window approach.

**Table S9. Climate distance thresholds by zone set.** Values are climate distance thresholds (CDT) for the normalized multivariate climate distance ($CDT_9$) or non-normalized climate variables (AHM to TD), except for N, which is the number of zones used to calculate CDT. CDTs were calculated for mid-elevation forested zones, and then the medians were averaged by ecological zone sets (E), geographic zone sets (G), Douglas-fir zone sets (PSME), and means for ecological and geographical zone sets (E + G).

| | Ecological (E) | | Geographic (G) | | | | Sum (N) or mean ($CDT_9$ to TD) | | | |
|---|---|---|---|---|---|---|---|---|---|---|
| Variable | BEC | EPA4 | CA | ID/MT | OR66 | WA66 | E | G | E + G | PSME |
| N* | 104 | 121 | 538 | 1184 | 403 | 316 | 225 | 2441 | 2666 | 166 |
| $CDT_9$† | 1.04 | 1.16 | 0.68 | 0.64 | 0.60 | 0.71 | 1.10 | 0.66 | 0.88 | 0.96 |
| AHM | 6.40 | 13.13 | 6.49 | 4.49 | 4.36 | 3.74 | 9.76 | 4.77 | 7.27 | 4.04 |
| CMD | 94.00 | 175.26 | 82.28 | 59.00 | 63.07 | 64.27 | 134.63 | 67.15 | 100.89 | 87.67 |
| DD_0 | 185.5 | 101.5 | 37.49 | 95.39 | 53.53 | 80.46 | 143.50 | 66.72 | 105.11 | 82.88 |
| DD5 | 262.44 | 535.28 | 292.45 | 149.04 | 187.52 | 172.18 | 398.86 | 200.30 | 299.58 | 244.34 |
| EMT | 2.96 | 4.10 | 3.15 | 1.70 | 1.91 | 1.98 | 3.53 | 2.19 | 2.86 | 2.74 |
| EREF | 78.75 | 153.76 | 73.00 | 45.94 | 51.50 | 46.33 | 116.25 | 54.19 | 85.22 | 69.59 |
| EXT | 2.25 | 3.05 | 1.35 | 1.05 | 1.00 | 1.13 | 2.65 | 1.13 | 1.89 | 1.64 |
| FFP | 19.5 | 34.00 | 26.50 | 14.50 | 18.72 | 17.00 | 26.75 | 19.18 | 22.96 | 24.25 |
| MAP | 261.94 | 369.00 | 286.17 | 172.58 | 248.47 | 464.03 | 315.47 | 292.82 | 304.14 | 703.35 |
| MAT | 1.47 | 2.20 | 1.10 | 0.80 | 0.75 | 0.86 | 1.84 | 0.88 | 1.36 | 1.15 |
| MCMT | 1.42 | 1.90 | 1.10 | 0.75 | 0.80 | 0.85 | 1.66 | 0.87 | 1.27 | 1.28 |
| MSP | 79.70 | 55.50 | 20.50 | 42.22 | 32.06 | 78.00 | 67.60 | 43.19 | 55.40 | 106.08 |
| MWMT | 1.82 | 2.50 | 1.45 | 0.95 | 0.99 | 1.00 | 2.16 | 1.10 | 1.63 | 1.27 |
| PAS | 173.80 | 91.50 | 33.04 | 114.46 | 70.50 | 134.07 | 132.65 | 88.02 | 110.33 | 169.91 |
| SHM | 16.30 | 63.95 | 66.78 | 11.10 | 18.77 | 16.36 | 40.12 | 28.25 | 34.19 | 21.58 |
| TD | 1.45 | 1.40 | 0.95 | 0.60 | 0.80 | 0.75 | 1.42 | 0.77 | 1.10 | 1.05 |

*N is the number of forested mid-elevation zones used in the analyses.

†CDTs were calculated using a nine-variable climate distance function or individual climate variables (AHM-TD). Climate variables are described in Table S2.

**Table S10. Climate distance thresholds (CDTs) inferred from provenance tests of eight conifer species.**

| Species | Climate Distance Threshold (CDT) |
|---|---|
| PICO | 0.807 |
| PIPO | 0.819 |
| PIMO | 0.875 |
| PSME | 0.948 |
| ABGR | 1.019 |
| THPL | 1.259 |
| TSHE | 1.291 |
| PIEN | 1.380 |

*See Table S1 for a description of species names from USDA plant symbols (USDA PLANTS database).

**Table S11. Seed deployment using the climate-based focal zone system, 9-variable climate distance function, current climate (1991-2020), and a climate distance threshold (CDT) of 0.9.** Numbers and areas of matching target zones (rows) are shown by focal zone set (columns).

| | Focal zone set | | | | | | | | | | | |
|---|---|---|---|---|---|---|---|---|---|---|---|---|
| | BEC | | EPA4 | | CA | | ID/MT | | OR66 | | WA66 | |
| | N | Area | N | Area | N | Area | N | Area | N | Area | N | Area |
| **(a) Zone set statistics** | | | | | | | | | | | | |
| Total* | 200 | 81.53 | 253 | 81.63 | 900 | 25.90 | 1807 | 34.54 | 678 | 15.44 | 555 | 13.33 |
| Zone mean† | 1 | 0.408 | 1 | 0.323 | 1 | 0.029 | 1 | 0.019 | 1 | 0.023 | 1 | 0.024 |
| **(b) Target zone matches in ecological zone sets** | | | | | | | | | | | | |
| BEC | 14.4 | 4.24 | 5.2 | 1.82 | 0.6 | 0.00 | 87.3 | 1.55 | 8.7 | 0.14 | 16.9 | 0.33 |
| EPA4 | 4.1 | 0.84 | 12.3 | 4.21 | 33.8 | 1.11 | 87.9 | 1.91 | 28.9 | 0.79 | 17.9 | 0.53 |
| **(c) Target zone matches in geographic zone sets** | | | | | | | | | | | | |
| CA | 0.1 | 0.01 | 9.5 | 3.03 | 78.6 | 1.99 | 3.8 | 0.09 | 26.8 | 0.75 | 5.5 | 0.14 |
| ID/MT | 9.7 | 2.21 | 12.3 | 4.46 | 1.9 | 0.04 | 192.8 | 3.92 | 13.7 | 0.28 | 21.1 | 0.47 |
| OR66 | 2.6 | 0.53 | 10.8 | 4.18 | 35.5 | 0.72 | 36.4 | 0.79 | 78.3 | 1.77 | 29.1 | 0.88 |
| WA66 | 6.1 | 1.55 | 8.2 | 3.19 | 9.0 | 0.15 | 68.7 | 1.43 | 35.5 | 0.68 | 47.1 | 1.18 |
| Total (unique)‡ | 25.70 | 6.81 | 31.75 | 11.36 | 80.00 | 2.01 | 282.10 | 5.62 | 100.30 | 2.28 | 77.25 | 1.93 |

*Total number (N) or area (Area, millions of ha) of forested zones.

†Zone mean is the zone number (N) or zone area divided by N (i.e., N/N or Area/N).

‡Because the EPA4 and geographic zone sets covered the same area in the U.S., these values were obtained by adding the BEC results to the mean of the results for EPA4 and the sum of the geographic zone sets.

### Figures

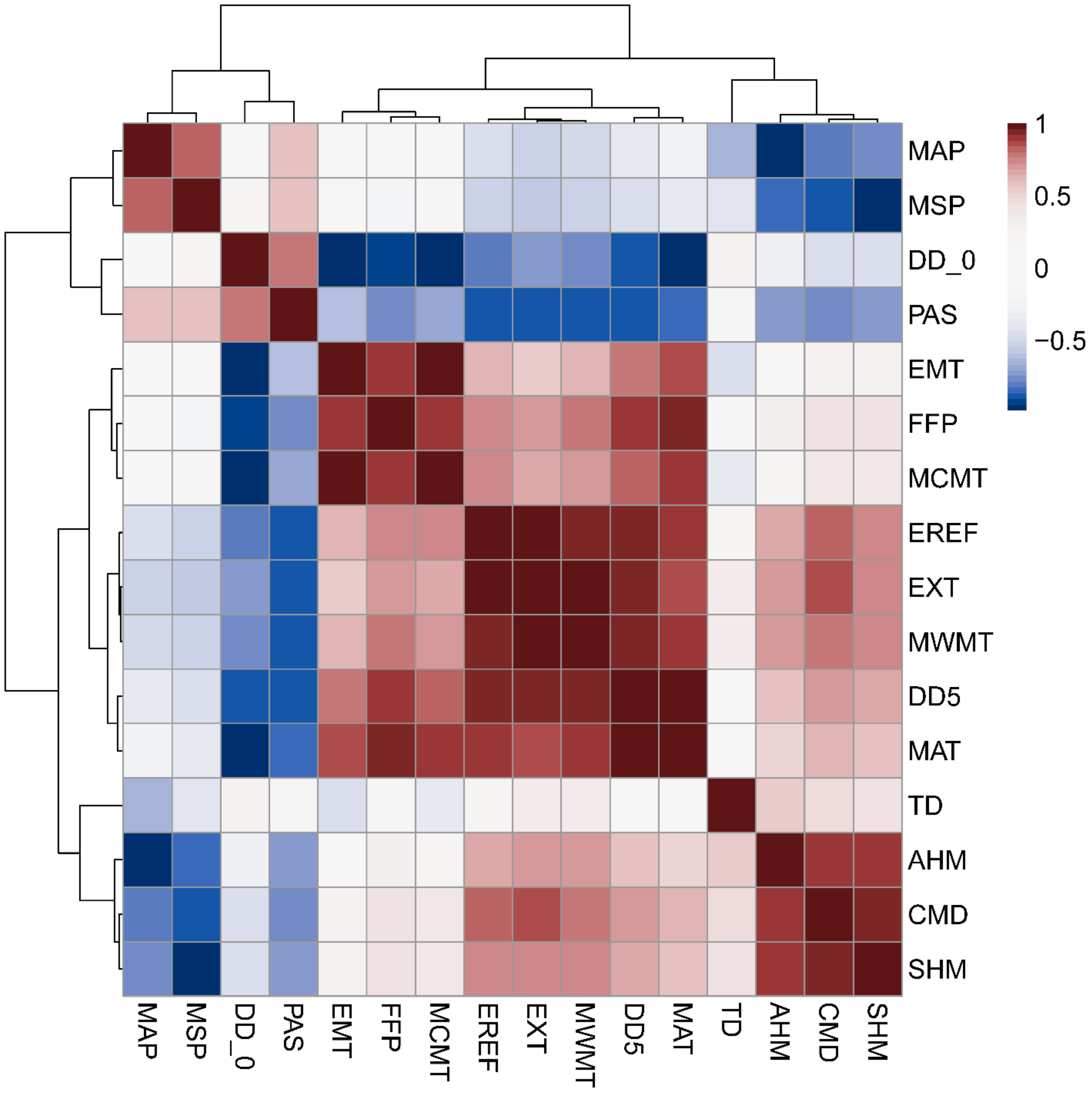


**Figure S1. Correlations of 16 climate variables for four non-overlapping Pacific Northwest geographic zone sets (CA, ID/MT, OR66, and WA66).** Values are averages of correlation coefficients from four zone sets. Climate variables are described in Table S2.

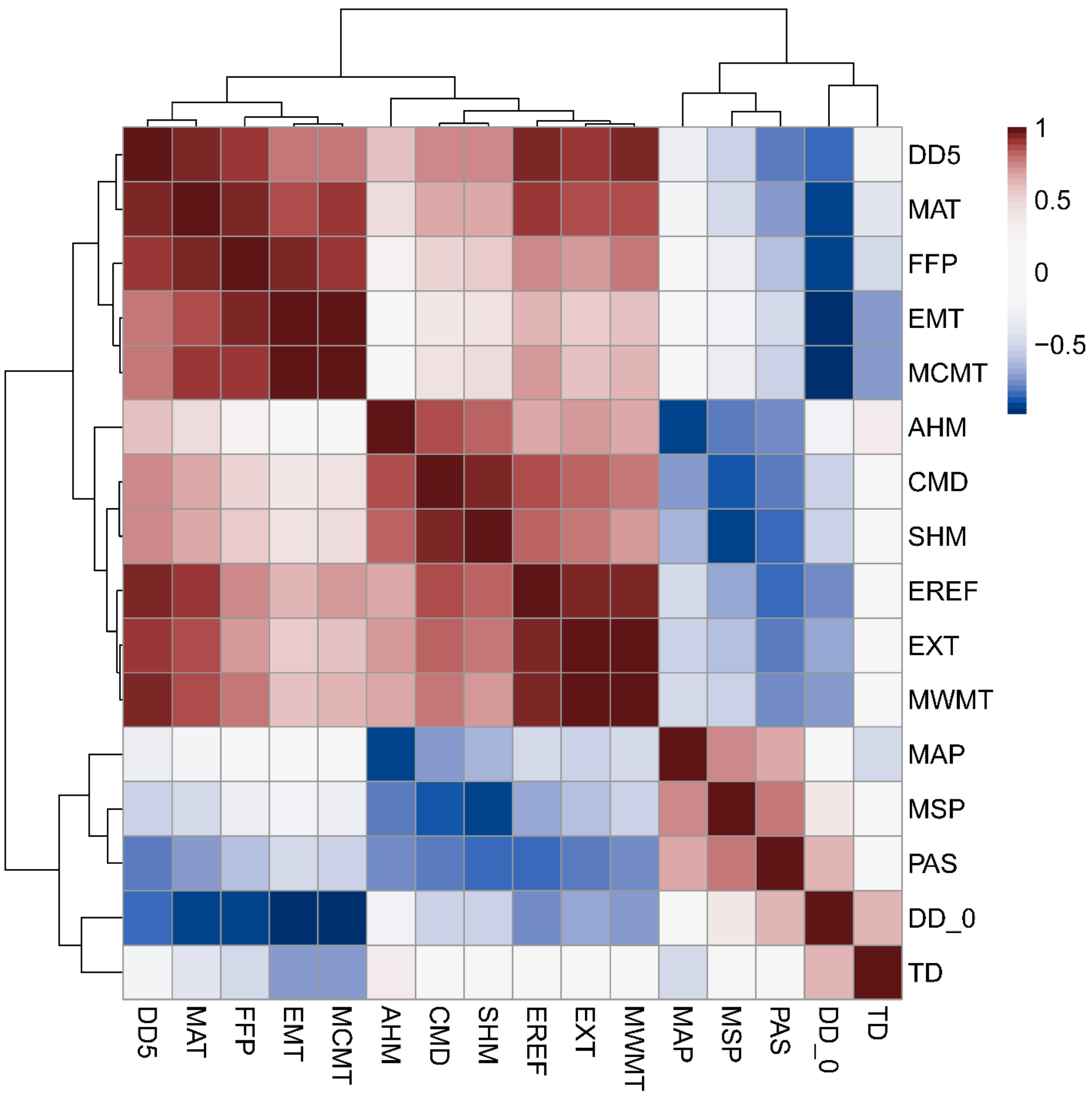


**Figure S2. Correlations of 16 climate variables for two non-overlapping Pacific Northwest ecological zone sets (BEC and EPA4).** Values are averages of correlation coefficients from two zone sets. Climate variables are described in Table S2.

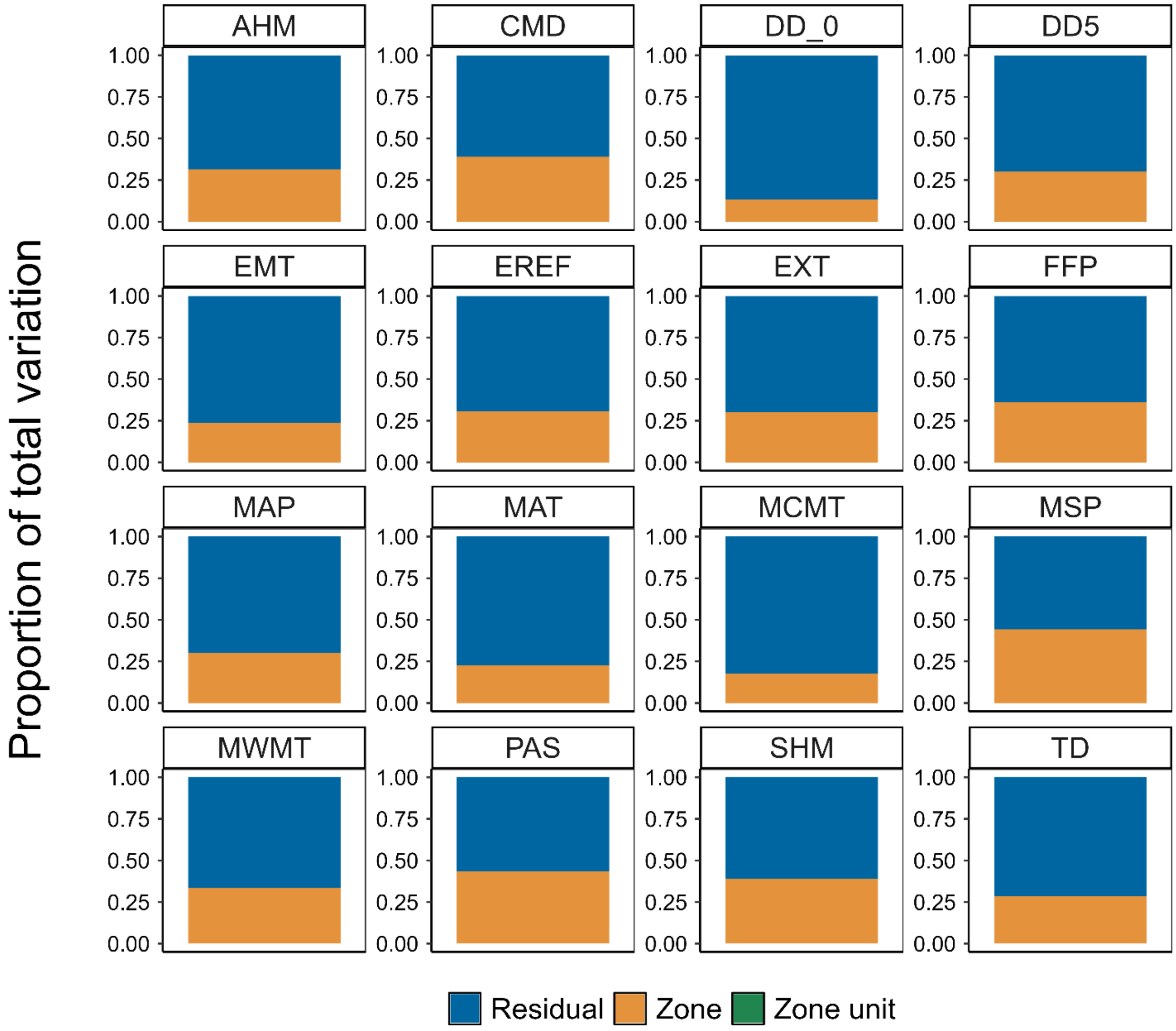


**Figure S3. Hierarchical partitioning of climate variation for one of two ecological zone sets (BEC).** In general, climate variation was partitioned into three components: within zones (Residual), among zones within zone units (Zone), and among zone units (Zone unit), and then expressed as a proportion of total variation. For the ecological zone sets, however, there was no distinction between zone units and zones. Abbreviations for ClimateNA variables are described in Table S2.

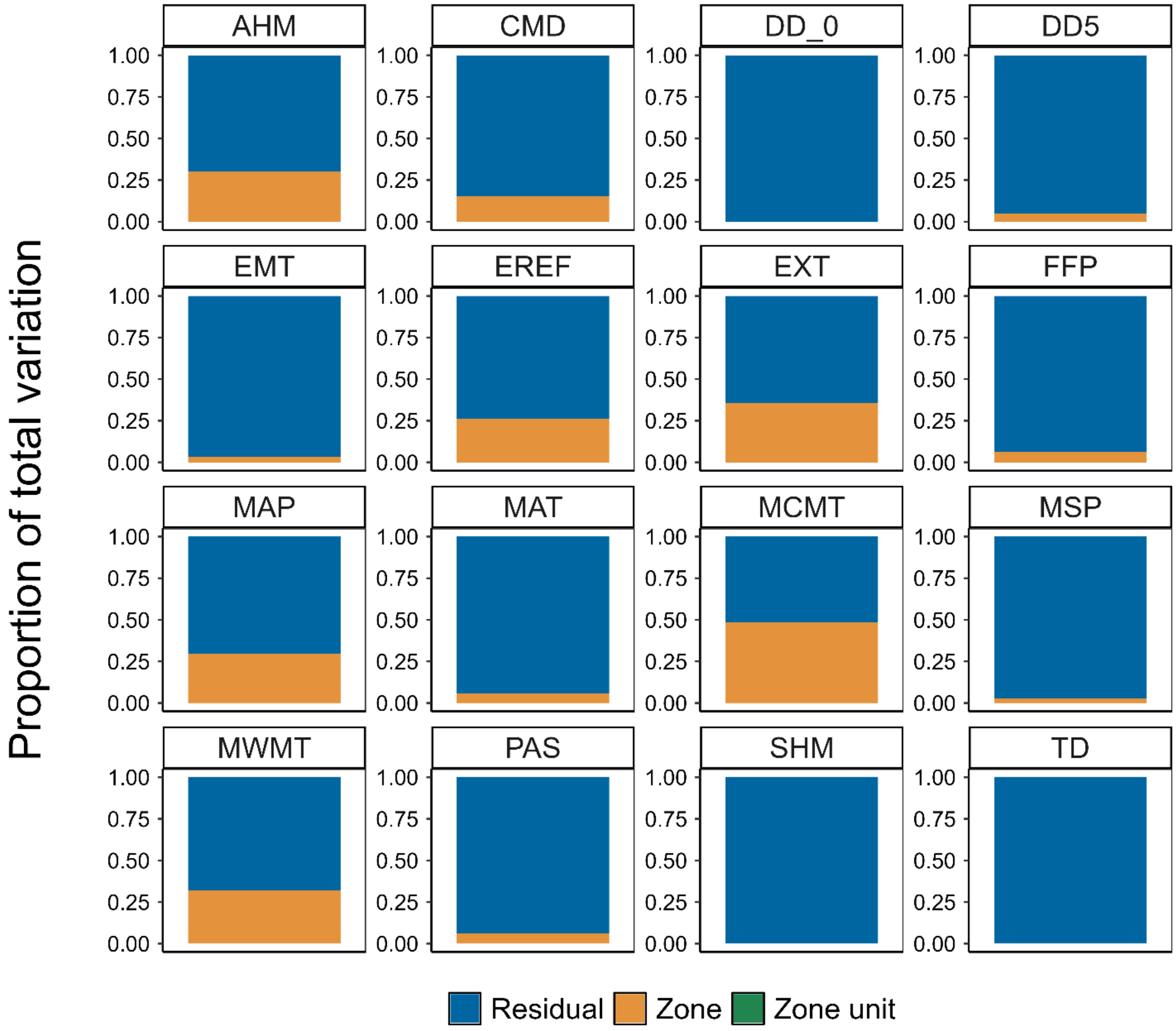


**Figure S4. Hierarchical partitioning of climate variation for one of two ecological zone sets (EPA4).** In general, climate variation was partitioned into three components: within zones (Residual), among zones within zone units (Zone), and among zone units (Zone unit), and then expressed as a proportion of total variation. For the ecological zone sets, however, there was no distinction between zone units and zones. Abbreviations for ClimateNA variables are described in Table S2.

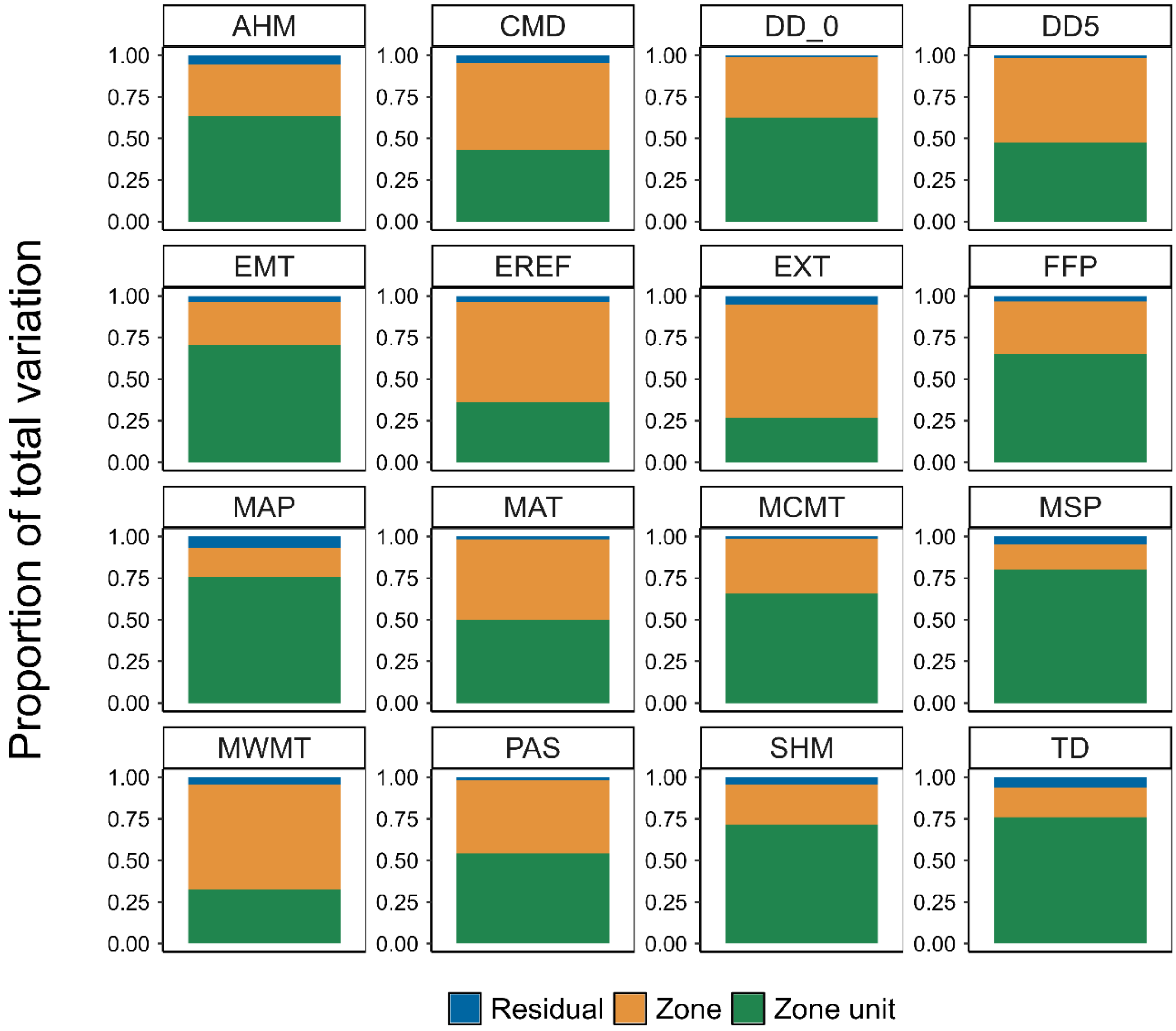


**Figure S5. Hierarchical partitioning of climate variation for one of four geographic zone sets (CA).** Climate variation was partitioned into three components: within zones (Residual), among zones within a zone unit (Zone), and among zone units (Zone unit), and then expressed as a proportion of total variation. Abbreviations for ClimateNA variables are described in Table S2.

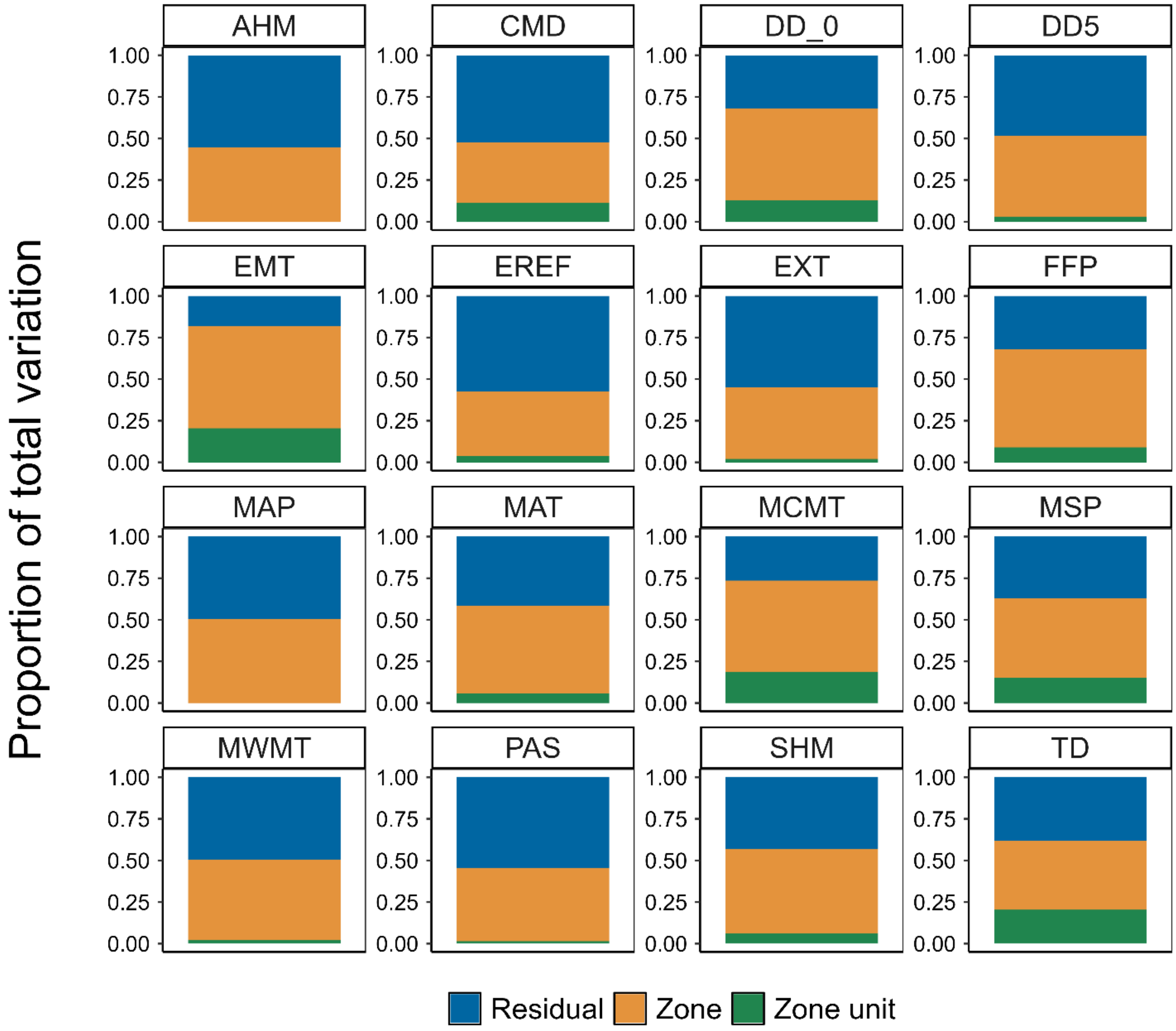


**Figure S6. Hierarchical partitioning of climate variation for one of four geographic zone sets (ID/MT).** Climate variation was partitioned into three components: within zones (Residual), among zones within a zone unit (Zone), and among zone units (Zone unit), and then expressed as a proportion of total variation. Abbreviations for ClimateNA variables are described in Table S2.

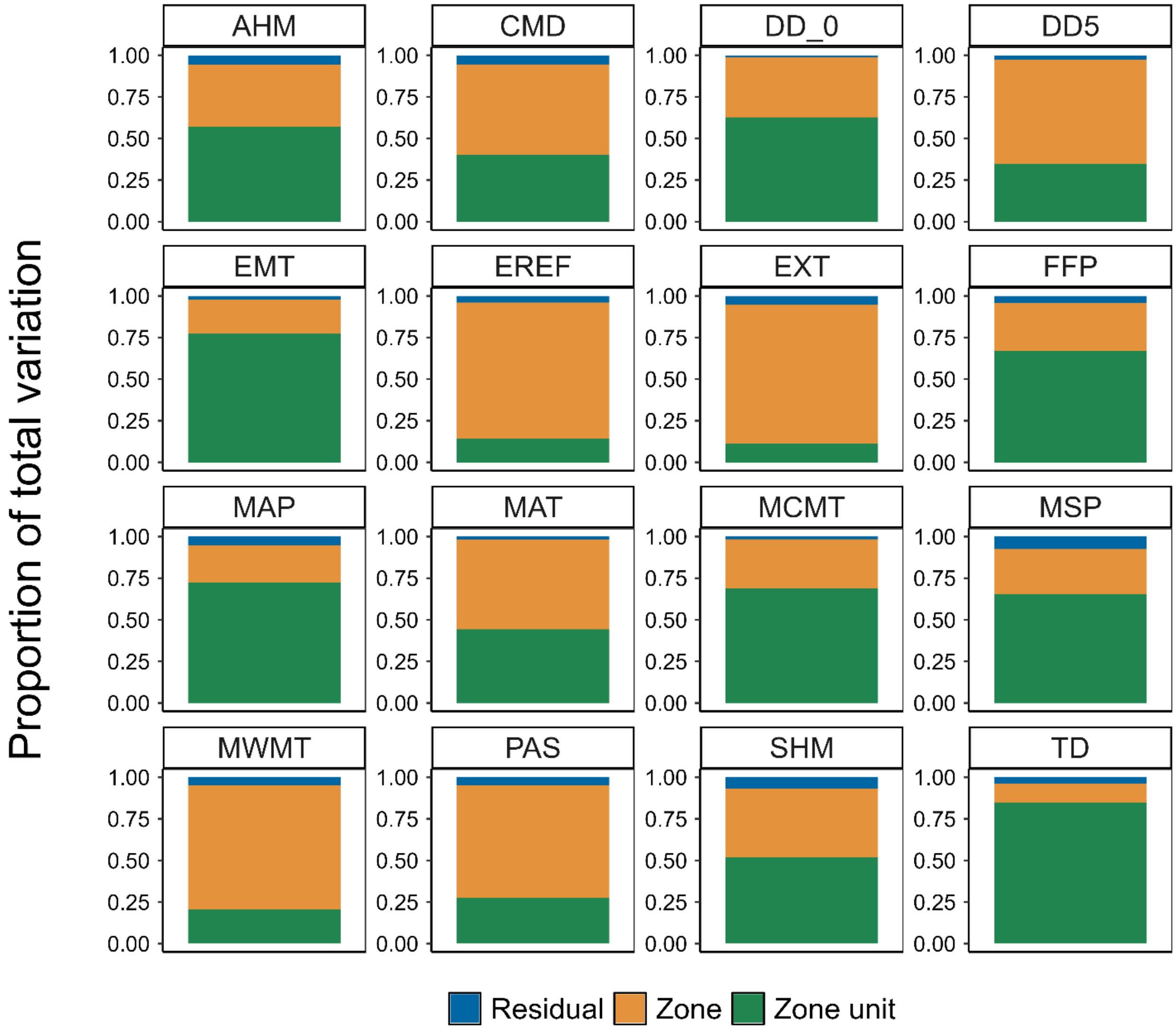


**Figure S7. Hierarchical partitioning of climate variation for one of four geographic zone sets (OR66).** Climate variation was partitioned into three components: within zones (Residual), among zones within a zone unit (Zone), and among zone units (Zone unit), and then expressed as a proportion of total variation. Abbreviations for ClimateNA variables are described in Table S2.

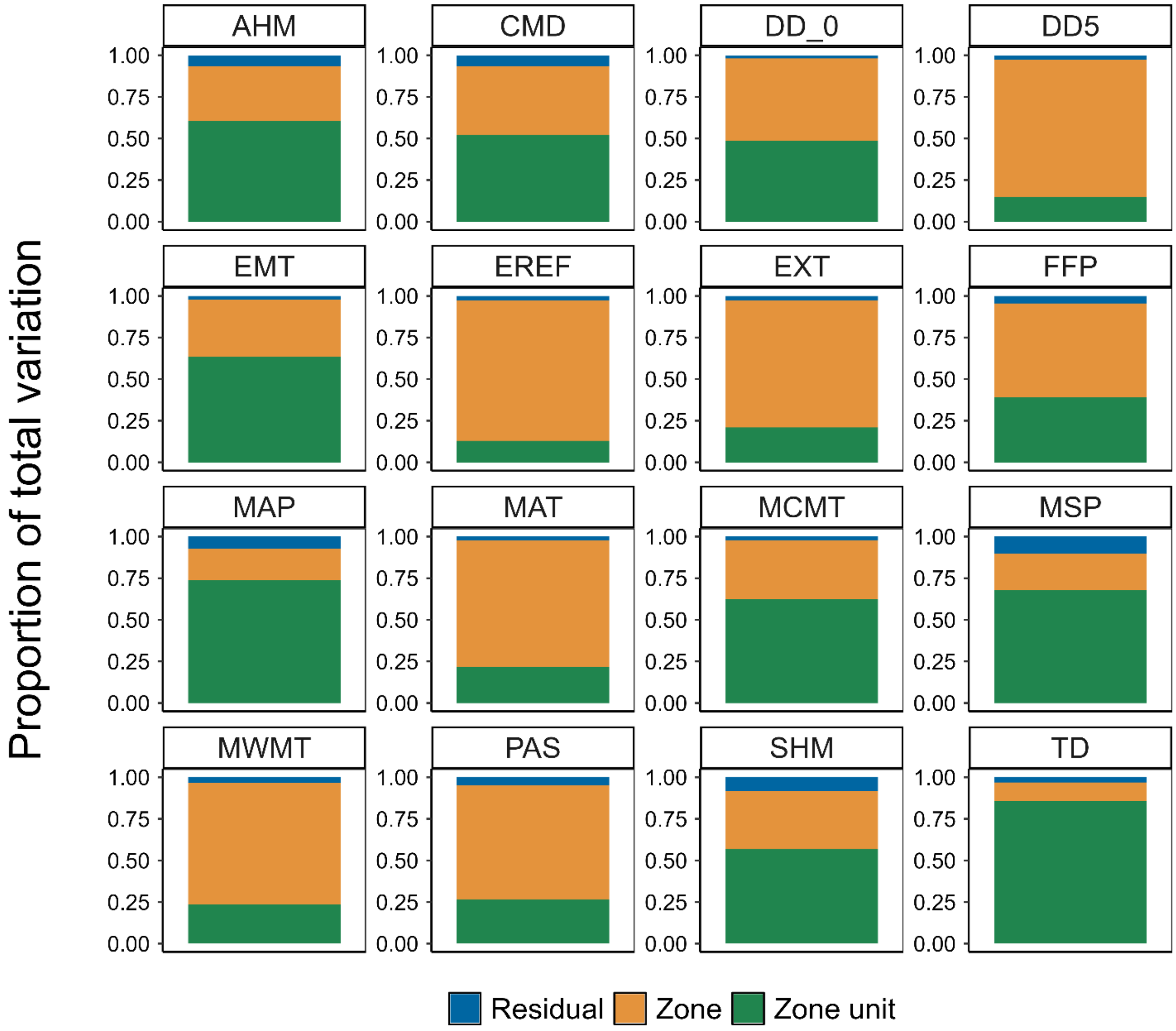


**Figure S8. Hierarchical partitioning of climate variation for one of four geographic zone sets (WA66).** Climate variation was partitioned into three components: within zones (Residual), among zones within a zone unit (Zone), and among zone units (Zone unit), and then expressed as a proportion of total variation. Abbreviations for ClimateNA variables are described in Table S2.

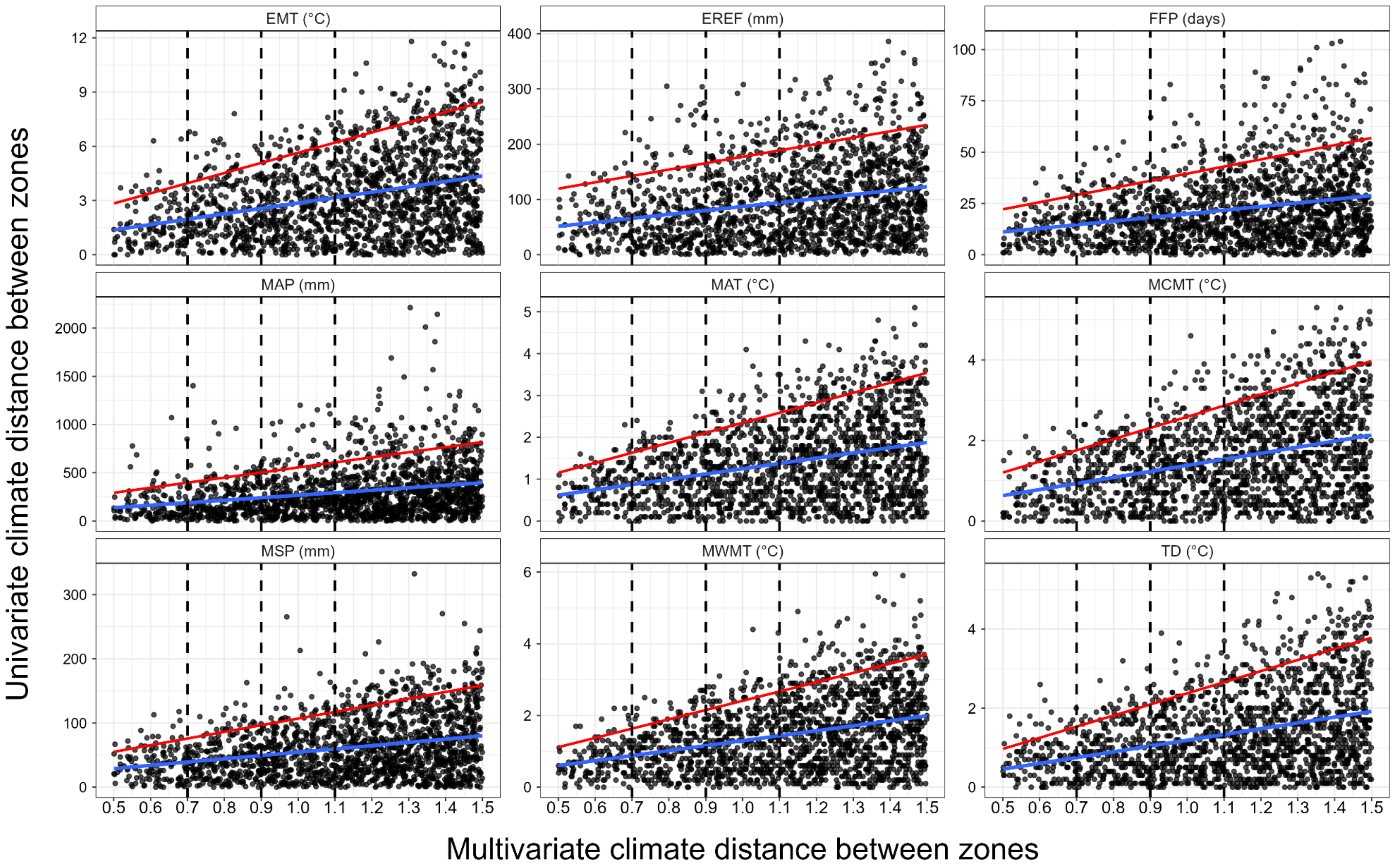


**Figure S9. Regression for predicting univariate climate distances (EMT-TD) from the multivariate (9-variable) climate distance.** Blue (lower) lines are smoothed conditional means from the regression, whereas red (upper) lines are the $90^{th}$ percentiles for individual predictions. Distances for MAT, MCMT, and MWMT rarely exceed 2°C when the Climate Distance Threshold (CDT) equals 0.9 (moderate). Horizontal lines also show the conservative (0.7) and liberal (1.1) CDTs used in Zone Matcher. Abbreviations for ClimateNA variables are described in Table S2.
